# Distinct Nrf2 Signaling Thresholds Mediate Lung Tumor Initiation and Progression

**DOI:** 10.1101/2022.08.24.504986

**Authors:** Janine M. DeBlasi, Aimee Falzone, Samantha Caldwell, Nicolas Prieto-Farigua, Justin R. Prigge, Edward E. Schmidt, Iok In Christine Chio, Florian A. Karreth, Gina M. DeNicola

## Abstract

Mutations in the KEAP1-NRF2 pathway occur in up to a third of non-small cell lung cancer (NSCLC) cases and often confer resistance to therapy and poor outcomes. Here, we developed murine alleles of the KEAP1 and NRF2 mutations found in human NSCLC and comprehensively interrogated their impact on tumor initiation and progression. Chronic Nrf2 stabilization by Keap1 or Nrf2 mutation was not sufficient to induce tumorigenesis, even in the absence of tumor suppressors p53 or Lkb1. When combined with Kras^G12D/+^, constitutive Nrf2 activation promoted lung tumor initiation and early progression of hyperplasia to low-grade tumors but impaired their progression to advanced-grade tumors, which was reversed by Nrf2 deletion. Finally, NRF2 overexpression in KEAP1 mutant NSCLC cell lines was detrimental to cell proliferation, viability, and anchorage-independent colony formation. Collectively, our results establish the context-dependence and activity threshold for NRF2 during the lung tumorigenic process.

**Significance:** This study reports murine lung cancer models harboring mutations in the Keap1/Nrf2 pathway and highlights the context-dependent and diverse roles of Nrf2 during lung tumor initiation and progression.

## Introduction

NRF2 (nuclear factor-erythroid 2 p45-related factor 2) is a stress-responsive transcription factor that regulates the detoxification of reactive oxygen species (ROS), maintains cellular homeostasis, and regulates many facets of metabolism (1, 2). NRF2 is negatively regulated by KEAP1 (Kelch-like ECH-associated protein 1), a substrate adaptor protein for the cullin 3 (CUL3)-based E3 ubiquitin ligase that facilitates NRF2 ubiquitination and proteasomal degradation in the absence of oxidative or xenobiotic stress (3). NRF2 promotes the detoxification of carcinogens to limit deleterious mutations that initiate cancer(4–7) and NRF2 activators are being explored as chemopreventative agents (8–10). However, NRF2 is frequently stabilized in many cancers, particularly non-small cell lung cancer (NSCLC), where mutations in the KEAP1-NRF2 pathway are found in up to 30% of cases (11, 12). NRF2 stabilization is associated with poor prognosis (13–20), resistance to chemo- and radiotherapy (21, 22), cancer cell survival (22), proliferation (23), metabolic reprogramming (20, 24–26), and metastasis (27). It remains unclear, however, whether chronic NRF2 stabilization transforms normal cells. Thus, it is important to understand the contexts and mechanisms by which NRF2 can prevent and promote cancer phenotypes.

Preclinical genetically engineered mouse models (GEMMs) have advanced our understanding of the role of NRF2 in lung tumorigenesis (23, 27–37). NRF2 activation in NSCLC has been modeled in GEMMs by inactivating Keap1 via conditional knockout (28-31, 33, 35) or CRISPR-mediated deletion (27, 34, 36, 37), in contrast to the KEAP1 mutations found in human lung cancer. These studies have shown that Nrf2 promotes lung tumor initiation (29, 30), tumor size (34, 38), progression (33, 34), and metastasis (27). However, other studies have failed to see an effect of Nrf2 on lung tumor initiation (38, 39) or size (36, 37, 39), and we reported that Nrf2 activation significantly decreases tumor size (32). Study conditions, time points and phenotypes assayed varied across these studies. Therefore, the role NRF2 activation plays at distinct stages of tumor initiation and progression remains to be determined.

In the current study, we generated lung cancer GEMMs expressing Keap1^R554Q^ and Nrf2^D29H^ mutations to comprehensively investigate how Nrf2 activation affects each stage of the tumorigenic process. These models also exhibit a series of graded NRF2 activation, allowing us to ask how different levels of Nrf2 influence lung tumor progression. We found that constitutive Nrf2 stabilization induced by these mutations was insufficient for lung tumor development, even in the context of tumor suppressor loss. In contrast, these mutations promoted lung tumor initiation in the Kras^G12D/+^ model of early lung adenocarcinoma, consistent with previous studies (23, 29, 30).

Using the Kras^G12D/+^; p53^fl/fl^ adenocarcinoma model, we found that homozygous Keap1 mutation unexpectedly blocked tumor progression. Supportingly, we found that Nrf2 expression and activity was downregulated in advanced tumors, and Nrf2 deletion could rescue the Keap1 mutation-mediated progression impairment. Overall, our data suggest that NRF2 has distinct, threshold-dependent effects during lung tumor initiation and progression.

## Materials and Methods

### Lead contact

Further information and requests for resources and reagents should be directed to and will be fulfilled by the Lead Contact, Dr. Gina M. DeNicola.

### Material availability

All unique reagents generated in this study are available from the Lead Contact without restriction.

### Mice

Mice were housed and bred in accordance with the ethical regulations and approval of the IACUC (protocols #: IS00003893R and IS00007922R). Generation of the Keap1 targeting vector was previously described (32). Briefly, the *CA-Keap1*^*R554Q*^ allele (Keap1^tm1Gmdn^, MGI: 7327097) was made by inserting a wild-type *Keap1* cDNA containing exons 3-5 flanked by loxP sites upstream of the R554Q mutation in exon 4 of the *Keap1* gene. *Keap1* was targeted in C10 murine ES cells and cells were selected with blasticidin. To make the *LSL-Nfe2l2*^*D29H*^ allele (Nfe2l2^tm1Gmdn^, MGI: 7327101), a STOP cassette flanked by loxP sites was inserted into intron 1 and codon 29 in endogenous exon 2 was mutated from an aspartic acid to a histidine. The endogenous *Nfe2l2* locus was targeted in C10 murine ES cells and puromycin was used to select positive cells. For both alleles, positive clones were screened by copy number real-time PCR and injected into blastocysts. Genotyping primers were as follows: for the *Keap1*^R554Q^ allele: Mutant forward: 5’-ATGGCCACACTTTTCTGGAC-3’; wild-type (WT) forward: 5’-GGGGGTAGAGGGAGGAGAAT-3’; Common reverse: 5’-GCCACCCTATTCACAGACCA-3’. The WT PCR product was 326 bp and the mutant PCR product 584 bp. For the *Nfe2l2*^D29H^ allele: WT forward: 5’-GAGGCAGGTAGTTCTCTGAGTTTG-3’; Common reverse: 5’-GCAAATGCACTGAGACACTCAT-3’; Mutant forward: 5’-CTAGCCACCATGGCTTGAGT-3’. The WT PCR product was 189 bp and the mutant PCR product 282 bp. All mice were maintained on a mixed C57BL/6 genetic background. In addition to Keap1^R554Q^ and Nrf2^D29H^ mice, p53^flox^ (RRID:IMSR_JAX:008462); Lkb1^flox^ (RRID:IMSR_JAX:014143); Nrf2^flox^ (RRID:IMSR_JAX:025433); and LSL-Kras^G12D/+^ (RRID:IMSR_JAX:008179) mice were used. For mouse lung tumor studies, intranasal installation of 2.5 × 10^7^ PFU adenoviral-Cre (University of Iowa) was used to induce lung tumors as previously described (40). Adenoviral infections were performed under isofluorane anesthesia.

### Murine embryonic fibroblast generation and culture

MEFs were isolated from E13.5-14.5-day old embryos and maintained in pyruvate-free DMEM (Corning) containing 10% FBS, 100 units/mL penicillin and 100 µg/mL streptomycin (Gibco) in a humidified incubator with 5% CO^2^ and 95% air at 37°C. MEFs were used within four passages and infected with control empty adenovirus or adenoviral-Cre (University of Iowa) at an approximate multiplicity of infection of 500.

### RNA-sequencing preparation and analysis

Samples were prepared using the RNeasy plus mini kit (Qiagen, 74134). RNA quality was checked with the QIAxcel RNA QC kit (Qiagen, 929104). Additional RNA QC, sequencing, mapping to the mouse genome, and analysis were performed by Novogene. Differentially expressed genes (DESeq2) with p < 0.05 were included in the volcano plot.

### Immunohistochemistry (IHC)

Mouse lung tissue was fixed with 10% formalin overnight, transferred to 70% ethanol and paraffin embedded to be sectioned. Unstained tissue sections were de-paraffinized in xylene followed by rehydration in a graded alcohol series. Antigen retrieval was performed by boiling in 10mM citrate buffer (pH 6). Antibodies used for IHC include affinity-purified NRF2 (1:150 or 1:300) (41), NQO1 (Sigma Aldrich, RRID:AB_1079501, 1:500), Ki-67 (Cell Signaling Technology, RRID:AB_2620142, 1:200), and Cleaved Caspase-3 (Cell Signaling Technology, RRID:AB_2070042, 1:1000). Following overnight incubation at 4°C in primary antibody, the ImmPRESS HRP goat anti-rabbit kit (Vector Laboratories, RRID:AB_2631198) was used as directed by manufacturer’s instructions. DAB peroxidase (HRP) substrate (Vector Laboratories, SK-4105) was used to develop immunohistochemical staining, followed by counterstaining with hematoxylin (Vector Laboratories, H-3404). Slides were scanned with the Aperio imager at 20x and the H-score of at least five representative regions/ mouse was analyzed with QuPath software(42). Representative images were captured using the Axio Lab.A1 microscope at 40x (Carl Zeiss Microimaging Inc.).

### Tumor grading analysis and histology

Lung tumor grading was performed manually as previously described (43). Tumor grading distribution percentages were calculated by dividing the number of tumors in a specific grade by the total number of tumors per mouse. Tumor burden by grade was calculated by dividing the area of the lung covered by a specific tumor grade by the total lung area.

### NSCLC cell lines and culture

Human lung cancer cell lines used include H1944 (RRID:CVCL_1508), H322 (RRID:CVCL_1556), A549 (RRID:CVCL_0023), HCC15 (RRID:CVCL_2057), and H460 (RRID:CVCL_0459) and were previously described (24). All cell lines were acquired from an authentic source (Hamon Cancer Center Collection). Cells were cultured in RPMI 1640 (Gibco) containing 5% FBS without antibiotics in a humidified incubator with 5% CO^2^ and 95% air at 37°C. Cells were confirmed to be free of mycoplasma with the MycoAlert kit (Lonza) immediately upon receipt and aliquots were frozen. Cell lines were tested monthly for mycoplasma and used within 10-20 passages.

### Lentivirus generation and infection of NSCLC cells

Lentiviruses were made by transfecting Lenti-X 293T cells (Takara 632180) overnight with polyethylenimine (PEI), lentiviral plasmid (pLX317-NRF2(44) or the control pLX317 empty vector(32)), and packaging plasmids pCMV-dR8.2 dvpr (RRID:Addgene_8455) and pCMV-VSV-G (RRID:Addgene_8454) in DMEM containing 10% FBS. To generate NRF2-overexpressing cells, NSCLC cells were transduced for 24 hrs with lentiviruses in medium containing polybrene (8 µg/mL). After transduction, infected cells were selected with 0.5 µg/mL (H1944, H322, H460) or 1 µg/mL (A549, HCC15) puromycin for 72 hrs. Immediately following selection, cells were seeded in respective puromycin concentrations for the indicated assays.

### Cell proliferation and cell death assays

NSCLC cells were monitored with the CELLCYTE X™ live cell imaging instrument (Cytena) over the course of 96 hours. Prior to imaging, SYTOX Green nucleic acid stain (Thermo Fisher Scientific, S7020) was added to medium at a final concentration of 20 nM. Images were acquired from each well at 8-hour intervals and analyzed using CellCyte Studio (CELLINK). Cell confluency was represented as the % of the image covered by cells. The number of dead cells was normalized to cell confluency [number of Sytox Green positive cells/ mm^2^/ cell confluency]. The area under the curve (AUC) values were calculated by summing the proliferation or normalized dead cell number at each time point.

### Western blotting

Cells were lysed in ice-cold RIPA buffer with protease inhibitors (Fisher Scientific, PIA32955) followed by sonication in a water bath sonicator (Diagenode). Protein was quantified using the DC protein assay (Bio-Rad). Lysates were prepared with 6X SDS sample buffer containing 12% (v/v) β-ME (VWR) and separated on Bolt™ or NuPAGE 4-12% Bis-Tris gels (Invitrogen). SDS-PAGE separation was followed by transfer to 0.45 µm nitrocellulose membranes (GE Healthcare). The membranes were blocked in 5% non-fat milk in Tris-buffered saline with 0.1% Tween 20 (TBST). For immunoblotting, the following antibodies diluted in 5% milk in TBST were used: KEAP1 (Millipore Sigma, RRID:AB_2921362, 1:2000), NRF2 (Cell Signaling Technologies, D1Z9C, RRID:AB_2715528, 1:1000), NQO1 (Sigma Aldrich, RRID:AB_1079501, 1:1000), GCLC (Sana Cruz Biotechnology, H-5, RRID:AB_2736837, 1:1000), xCT (abcam, RRID:AB_778944, 1:1000), GSR (Santa Cruz Biotechnology, RRID:AB_2295121, 1:1000), β-actin (Invitrogen AM4302, RRID:AB_2536382, 1:100,000). HRP secondary antibodies used include goat anti-rabbit IgG (Jackson ImmunoResearch Labs, RRID: AB_2313567), goat anti-mouse IgG (Jackson ImmunoResearch Labs, RRID:AB_10015289), and goat anti-rat IgG (Jackson ImmunoResearch Labs, RRID:AB_2338128). Membranes were developed with Clarity ECL substrate (Bio-Rad) or a luminol-based homemade ECL substrate.

### Soft agar colony formation assays

6-well plates were coated with a 0.8% agar prepared in RPMI. NSCLC cells were then seeded in 0.4% agar in RPMI. After the cell/ agar mixture solidified, RPMI medium containing 10% FBS, Pen/Strep and puromycin was added to each well and replenished every few days. Colonies were allowed to form for 10–16 days, and wells were stained with 0.01% crystal violet in a 4% paraformaldehyde in PBS solution. Plates were scanned on a flatbed scanner and ImageJ was used to quantify colonies.

### DepMap Analysis

*NFE2L2* dependency scores were downloaded from the DepMap database v. 22Q2 (45). Values were plotted from CRISPR (DepMap 22Q2 Public+Score, Chronos) for non-small cell lung cancer cell lines that we previously evaluated for high or low NRF2 activity (24).

### Quantification and statistical analysis

Graphpad Prism9 software was used for statistical analyses and P values < 0.05 were considered significant, with symbols as follows: *p<0.05, **p<0.01, ***p<0.001, ****p<0.0001. All data is represented as mean +/− standard deviation unless otherwise stated. For all experiments, similar variances between groups were observed.

### Data Availability

The data generated in this study are included in the article and its supplementary figures. Raw data are available upon request without restriction from the corresponding author. The gene expression data generated in this study are publicly available in Gene Expression Omnibus (GEO) at GSE212942.

## Results

### *Keap1*^*R554Q*^ and *Nfe2l2*^*D29H*^ alleles activate the Nrf2 transcriptional program

To study the role of NRF2 activation in lung cancer, we developed alleles harboring either the Keap1^R554Q^ or the Nrf2^D29H^ mutation found in human NSCLC **(Fig. 1A** and **B)**. Both Keap1^R554Q^ and Nrf2^D29H^ mutations prevent Keap1-mediated ubiquitination of Nrf2, allowing for constitutive expression of Nrf2 and transcription of Nrf2 target genes (46, 47). To generate the conditionally active *(CA)-Keap1*^*R554Q*^ allele, we inserted a wild-type *Keap1* cDNA containing exons 3-5 flanked by loxP sites upstream of the R554Q mutation in endogenous exon 4 of the *Keap1* gene **(Fig. 1A)** (32). For the Lox-STOP-Lox *(LSL)-Nfe2l2*^*D29H*^ allele, we inserted a loxP-flanked transcriptional and translational STOP (LSL) cassette upstream of the D29H mutation in exon 2 of the endogenous *Nfe2l2* gene **(Fig. 1B)**. For both alleles, Cre-mediated excision of loxP-flanked cassettes allows for physiological expression of Keap1^R554Q^ or Nrf2^D29H^, recapitulating NRF2 activation in human NSCLC. To validate the functionality of these alleles, we first generated mouse embryonic fibroblasts (MEFs), which allowed the switching from a Nrf2 deficient state (Nrf2^LSL/LSL^) to a Nrf2 stabilized state (Nrf2^D29H/D29H^), or from a basal Nrf2 state (Keap1^+/+^) to a stabilized Nrf2 state (Keap1^R554Q/R554Q^) in an isogenic system. Using these MEFs, we performed both transcriptomic profiling **(Fig. 1C** and **D)**. RNA-sequencing indicated that both Keap1^R554Q/R554Q^ and Nrf2^D29H/D29H^ MEFs demonstrated increased transcription of canonical Nrf2 target genes, including Nqo1, Srxn1, Txnrd1, and Gclc **(Fig. 1C** and **D)**. Prior targeting of the murine *Keap1* locus to generate a *Keap1*^*flox*^ allele resulted in the generation of a hypomorphic allele prior to Cre-mediated recombination, leading to decreased Keap1 levels and increased Nrf2 transcriptional activity throughout the whole animal (48). Importantly, we found no differences in expression of Keap1, Nrf2, or Nrf2 target proteins Nqo1 and Gclc between CA-Keap1^R554Q^ and WT Keap1 MEFs, indicating that the *CA-Keap1*^R554Q^ allele is not hypomorphic (Supplementary Fig. S1). Collectively, these results indicate that the mutant *Keap1*^*R554Q*^ and *Nfe2l2*^*D29H*^ alleles activate the Nrf2 transcriptional program.

**Figure 1.**
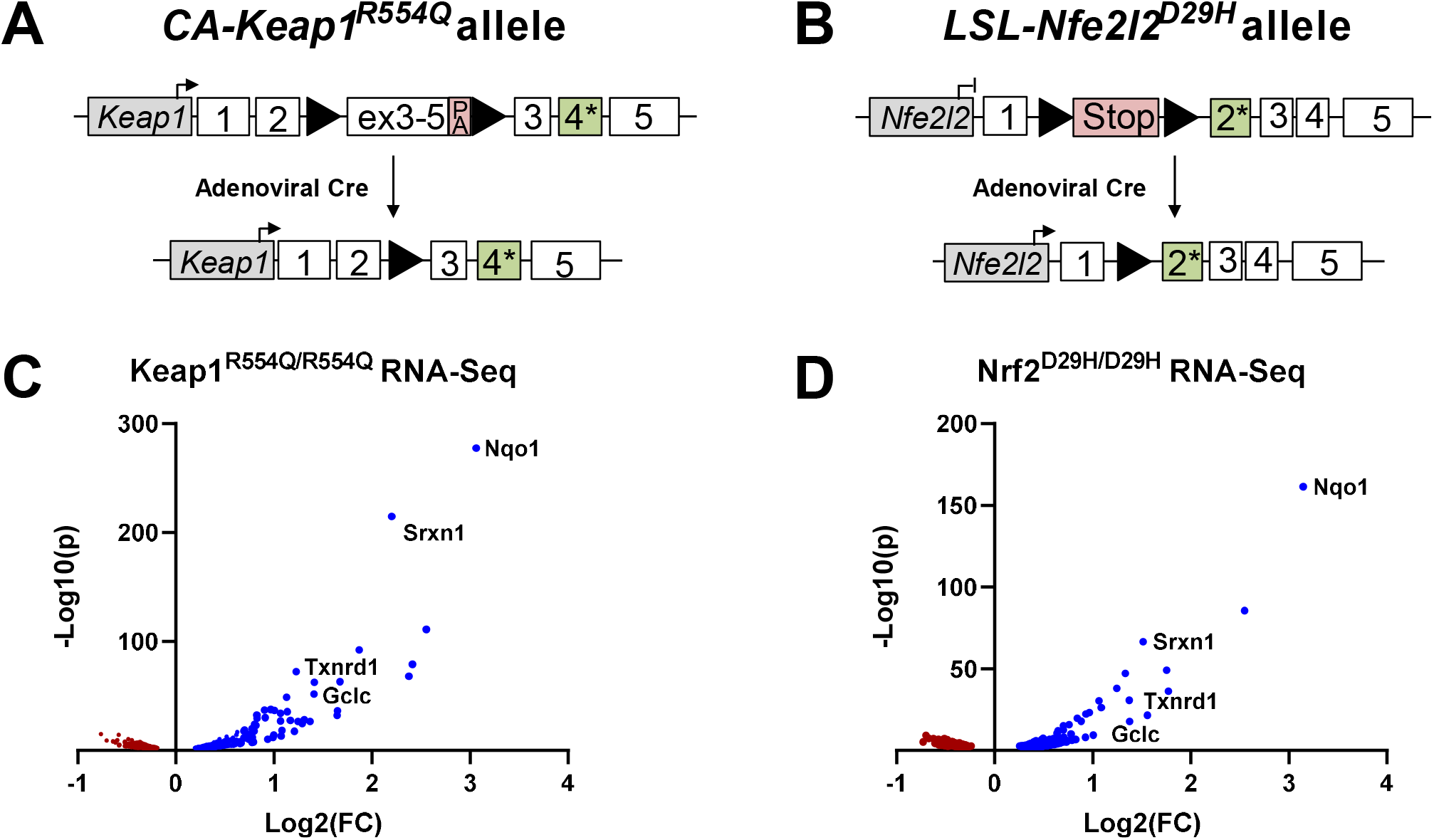
Development of mutant *Keap1* and *Nfe2l2* alleles found in human lung cancer. **A**, The conditionally active (CA)-*Keap1*^*R554Q*^ allele was generated by inserting a loxP-flanked, wild type (WT) *Keap1* cDNA containing exons 3-5 into intron 2 and introducing the R554Q mutation into endogenous exon 4 of the *Keap1* gene. Prior to intranasal installation of adenoviral-Cre recombinase Keap1 is wild type. Once the floxed cargo is excised, mutant Keap1^R554Q^ is expressed. PA = poly A signal. **B**, The Lox-STOP-Lox *(LSL)-Nfe2l2*^*D29H*^ allele was created by inserting a STOP cassette flanked by loxP sites into intron 1 and introducing the D29H mutation in endogenous exon 2 of the *Nfe2l2* gene. Following Cre-mediated excision of the STOP cassette, mutant Nrf2^D29H^ is expressed. **C**, Volcano plot of RNA-sequencing data from murine embryonic fibroblasts (MEFs) expressing Keap1^R554Q/R554Q^ compared to Keap1^+/+^. N=3, representative of two individual MEF lines. **D**, Volcano plot of RNA-sequencing data from MEFs expressing Nrf2^D29H/D29H^ compared to Nrf2^LSL/LSL^, which lack Nrf2 expression. N=3, representative of two individual MEF lines.

### Keap1 or Nrf2 mutation is not sufficient to initiate lung tumorigenesis

Given the importance of NRF2 in cytoprotection and redox homeostasis, there has been a longstanding interest in activating NRF2 pharmacologically for chemoprevention(8–10). The long-term safety of this approach, and whether the chronic activation of NRF2 can transform healthy cells *in vivo*, remains unknown. Moreover, the whole body deletion of Keap1 in mice results in postnatal lethality because of constitutive Nrf2 activation (49). In human lung tumors, *KEAP1* inactivation is frequently biallelic(13), whereas *NFE2L2* mutations are frequently heterozygous (50). Therefore, we induced the recombination of *Keap1* and *Nfe2l2* alleles in the lungs of Keap1^R554Q/+^, Keap1^R554Q/R554Q^, or Nrf2^D29H/+^ mice using adenoviral-Cre to test whether constitutive Nrf2 activation is sufficient to initiate lung tumor formation **(Fig. 2)**. First, we analyzed the overall survival between the different groups. The median survival across genotypes ranged from approximately 650-750 days, with no significant survival differences observed between wild-type and Keap1/Nrf2 mutant groups **(Fig. 2A)**. While mice did develop tumors, they comprised age-associated tumors like lymphoma. Upon examination of mouse lung histology for the presence of lung tumors, lung tumor-free survival was also not different between the groups **(Fig. 2B)**. Finally, histological analysis of lung tissues revealed that both alveolar and bronchiolar cells appeared phenotypically normal across the genotypes **(Fig. 2C)**. These results indicate that constitutive Nrf2 activation is not sufficient to induce lung tumor formation.

**Figure 2.**
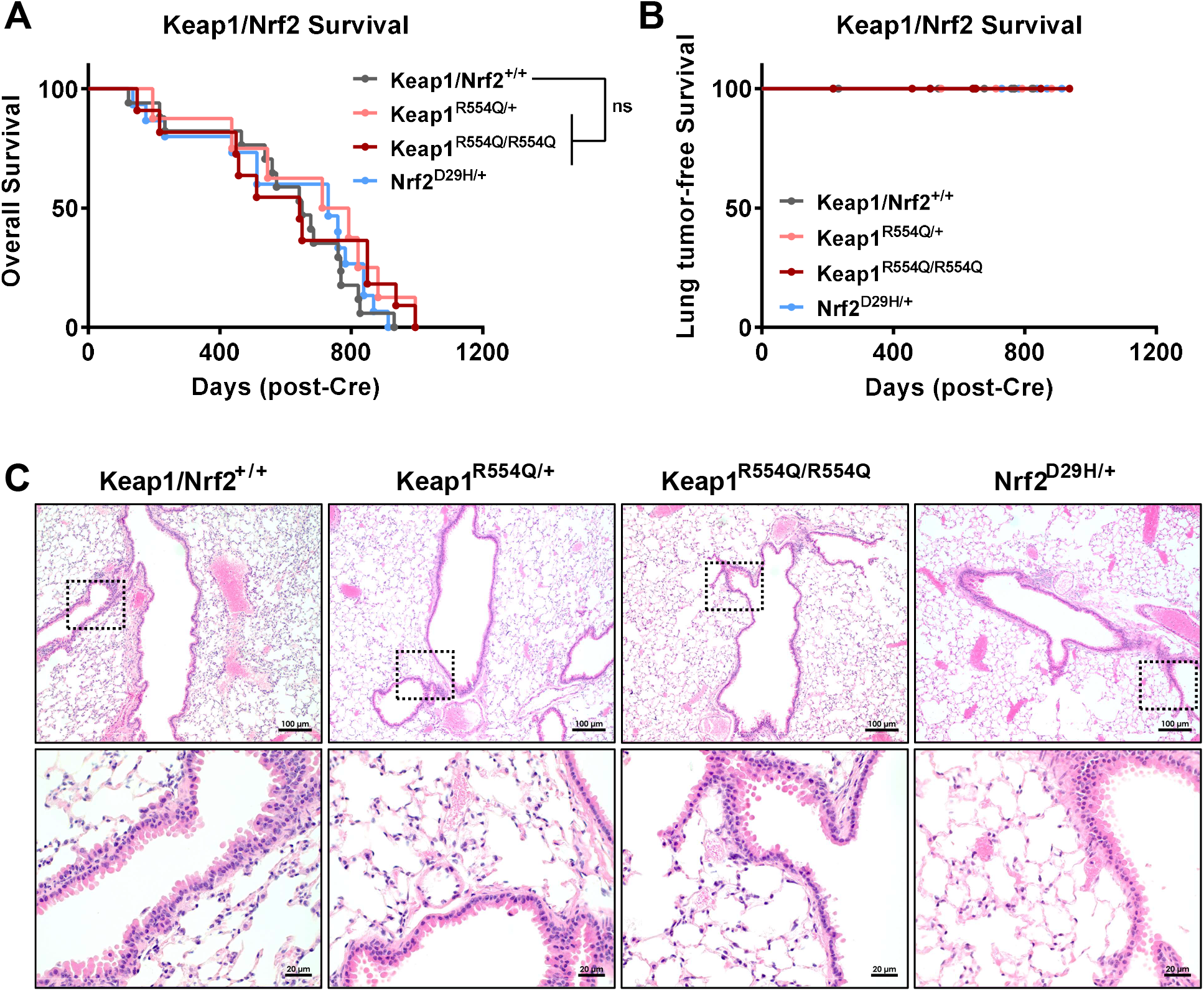
Mutation of Keap1 or Nrf2 is not sufficient to initiate lung tumorigenesis. **A**, Overall survival of Keap1/Nrf2 mutant mice. Keap1/Nrf2^+/+^ (n=17), Keap1^R554Q/+^ (n=8); Keap1^R554Q/R554Q^ (n=11); Nrf2^D29H/+^ (n=15). Ns= not significant (Log-rank (Mantel-Cox) test). **B**, Lung tumor-free survival of Keap1/Nrf2 mutant mice. Keap1/Nrf2^+/+^ (n=11), Keap1^R554Q/+^ (n=4); Keap1^R554Q/R554Q^ (n=8); Nrf2^D29H/+^ (n=11). **C**, Representative H&E of mouse lung depicting normal bronchiolar and alveolar cells (scale bars = 100μM (top panel), 20 μM (bottom panel)).

### Keap1 or Nrf2 mutation is not sufficient to initiate lung tumorigenesis in combination with tumor suppressor loss

To determine whether tumor suppressor loss was required for mutant Keap1 or Nrf2 to initiate lung tumor formation, we crossed Keap1 and Nrf2 mutant mice with p53^flox^ and Lkb1^flox^ alleles to concomitantly activate Nrf2 and delete these tumor suppressors in the lung **(Fig. 3)**. We first examined the consequence of Nrf2^D29H/+^, Keap1^R554Q/+^, or Keap1^R554Q/R554Q^ in combination with p53 deletion. Mice were aged to 500 days, at which time all mice were euthanized and examined for evidence of lung tumor formation. While a small number of these mice did succumb to disease prior to 500 days, they developed age-associated tumors including lymphoma, and we did not observe any differences in overall or lung tumor-free survival between genotypes **(Fig. 3A** and **B**). However, when examining the lung tissue histology, we observed dysplasia in Keap1^R554Q/R554Q^ bronchioles following loss of p53 **(Fig. 3C)**. This observation is consistent with previous work showing that tracheospheres derived from *Keap1*^*−/−*^;*Trp53*^*−/−*^ cells had an aberrant morphology (31). We next examined the consequence of Nrf2^D29H/+^, Keap1^R554Q/+^ or Keap1^R554Q/R554Q^ in combination with Lkb1 deletion. Similar to what was observed with p53, we also did not find any differences in overall or lung tumor-free survival between cohorts **(Fig. 3D** and **E)**. Moreover, the bronchiolar and alveolar morphology was normal across genotypes, in contrast to what was observed upon p53 loss **(Fig. 3F)**. Our findings indicate that Keap1/Nrf2 mutation is not sufficient to initiate lung tumor formation in combination with tumor suppressor loss.

**Figure 3.**
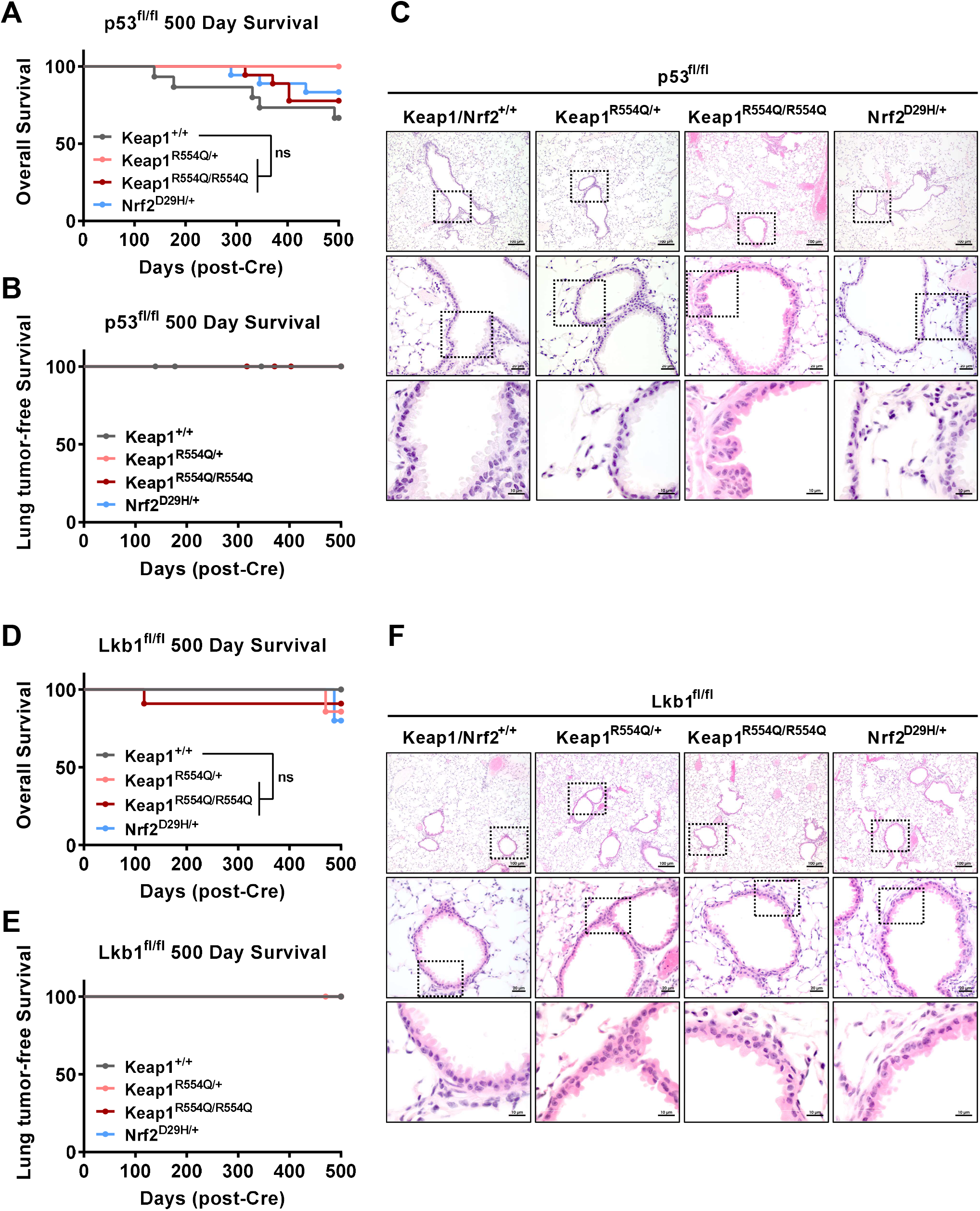
Mutation of Keap1 or Nrf2 is not sufficient to initiate lung tumorigenesis with p53 or Lkb1 loss. **A**, Overall survival of p53^fl/fl^ mice expressing wild-type or mutant Keap1/Nrf2. Keap1/Nrf2^*+/+*^ (n=15); Keap1^R554Q/+^ (n=7); Keap1^R554Q/R554Q^ (n=18); Nrf2^D29H/+^ (n=18). Ns= not significant (Log-rank (Mantel-Cox) test). **B**, Lung tumor-free survival of p53^fl/fl^ mice expressing wild-type or mutant Keap1/Nrf2. Keap1/Nrf2^+/+^ (n=11); Keap1^R554Q/+^ (n=6); Keap1^R554Q/R554Q^ (n=16); Nrf2^D29H/+^ (n=10). **C**. Representative H&E of mouse lung depicting bronchiolar and alveolar cells of the p53^fl/fl^ models (scale bars = 100μM (top panel), 20μM (middle panel), 10μM (bottom panel)). **D**, Overall survival of Lkb1^fl/fl^ mice expressing wild-type or mutant Keap1/Nrf2. Keap1/Nrf2^+/+^ (n=11); Keap1^R554Q/+^ (n=7); Keap1^R554Q/R554Q^ (n=11); Nrf2^D29H/+^ (n=5). Ns= not significant (Log-rank (Mantel-Cox) test). **E**, Lung tumor-free survival of Lkb1^fl/fl^ mice expressing wild-type or mutant Keap1/Nrf2. Keap1/Nrf2^+/+^ (n=11); Keap1^R554Q/+^ (n=6); Keap1^R554Q/R554Q^ (n=9); Nrf2^D29H/+^ (n=4). **F**, Representative H&E of mouse lung depicting bronchiolar and alveolar cells of the Lkb1^fl/fl^ models (scale bars = 100μM (top panel), 20μM (middle panel), 10μM (bottom panel)). For **A, B, D, E**, mice were infected intranasally with adenoviral-Cre, followed by collection at 500 days to analyze lung tissue histology.

### Nrf2 activation cooperates with mutant Kras to promote lung tumor initiation and early progression

We and others have reported that Nrf2 activation is important for Kras mutant lung tumorigenesis (23, 29, 30). To understand how Keap1 or Nrf2 mutation affect lung tumor initiation and early progression, we crossed Keap1/Nrf2 mutant mice with the Kras^G12D/+^ model of early lung adenocarcinoma (40). While all mice succumbed to lung tumors with a median survival of around 200 days, we observed no difference in survival between cohorts **(Fig. 4A** and **B)**. We next validated that these mutations were activating toward Nrf2 in tumors by performing immunohistochemical staining for Nrf2 and the Nrf2 target Nqo1. We observed that Keap1^R554Q/R554Q^ expression resulted in the greatest degree of Nrf2 activation, followed by Nrf2^D29H/+^, and then Keap1^R554Q/+^ compared to Keap1/Nrf2^+/+^ expression **(Fig. 4C-F)**. To examine the influence of Nrf2 activation on tumor initiation, we quantified tumor number across the genotypes and found that Keap1^R554Q/R554Q^ and Nrf2^D29H/+^ significantly increased tumor number in the Kras^G12D/+^ model **(Fig. 4G** and **H)**, consistent with prior reports using Keap1 deletion models (29, 30). We then examined the influence of Nrf2 on tumor progression by analyzing tumor grade. The distribution of atypical adenomatous and bronchiolar hyperplasia (AAH and BH, respectively) and tumors from grades 1 (adenoma) to 5 (adenocarcinoma) was determined. We observed an increase in the proportion and number of grade 1 tumors in Keap1^R554Q/R554Q^ and Nrf2^D29H/+^ mice compared to Keap1^R554Q/+^ and Keap1/Nrf2^+/+^ mice **(Fig. 4I**, Supplementary Fig. S2A). We also examined tumor size by grade and found that the median tumor size did not differ across genotypes (Supplementary Fig. S2B), although there were some modest differences across the individual grades, including decreased AAH size in all Keap1/Nrf2 mutant models (Supplementary Fig. S2C). Moreover, we found that there was a decrease in grade 3 tumor burden across all Keap1 and Nrf2 mutant models **(Fig. 4J)**, which resulted in a decreased overall tumor burden (Supplementary Fig. S2D) because grade 3 tumors were much larger than other grades (Supplementary Fig. S2C). Grade 3 tumors were extremely rare in the Kras^G12D/+^ model, however, limiting our ability to draw conclusions on the influence of Nrf2 activation on a limited number of tumors. Our findings indicate that Kras^G12D/+^ mutation cooperates with Keap1/Nrf2 mutation to promote formation of lung tumors and early progression to low-grade tumors.

**Figure 4.**
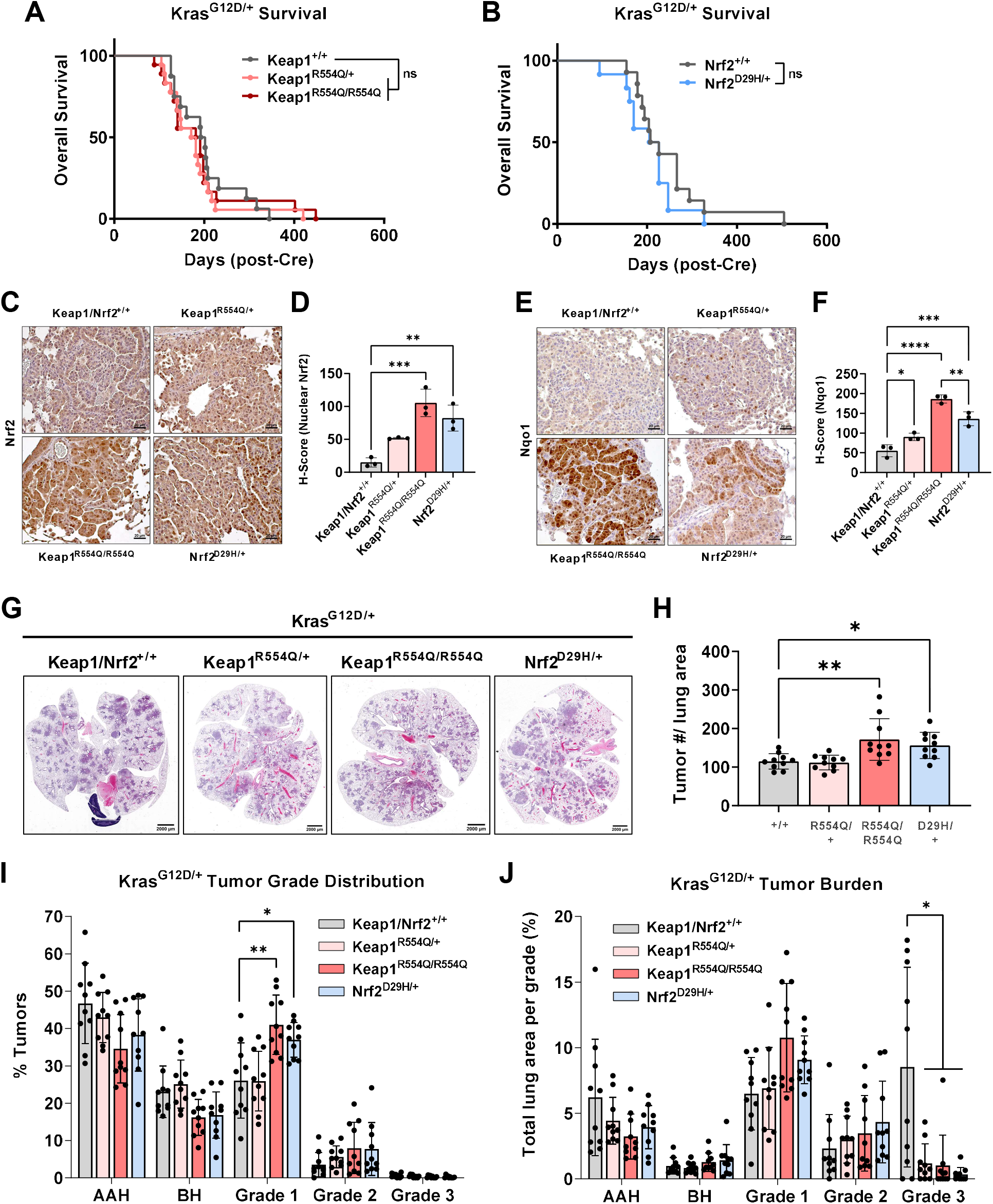
Keap1/Nrf2 mutation cooperates with Kras^G12D/+^ to promote lung tumor initiation and early progression. **A**, Overall survival of Keap1 mutant mice with Kras^G12D/+^ mutation. Keap1^+/+^ (n=16); Keap1^R554Q/+^ (n=18); Keap1^R554Q/R554Q^ (n=18). **B**, Overall survival of Nrf2 mutant mice with Kras^G12D/+^ mutation. Nrf2^+/+^ (n=14); Nrf2^D29H/+^ (n=12). Ns = not significant (Log-rank (Mantel-Cox) test). **C**, Representative immunohistochemical (IHC) staining of Nrf2 in Keap1/Nrf2 mutant tumors with Kras^G12D/+^ mutation (scale bars = 20 μM). **D**, H-scores for Nrf2 (nuclear) IHC staining. **E**, Representative immunohistochemical (IHC) staining of Nrf2 target Nqo1 (scale bars = 20 μM). **F**, H-scores for Nqo1 (whole cell) IHC staining. For **C-F**, N=3 mice per genotype and >20,000 tumor cells per mouse. *p<0.05 (one-way ANOVA). **G**, Representative whole lung H&E-stained section (scale bars = 2000 μM). **H**, Tumor number per mouse in Keap1/Nrf2 mutant models normalized to lung area. *p<0.05 (one-way ANOVA). **I**, Distribution of tumor grades across Keap1/Nrf2 mutant models. *p<0.05 (unpaired t test with Holm-Sidak’s multiple comparisons test). AAH = atypical adenomatous hyperplasia. BH = bronchiolar hyperplasia. **J**, Fraction of lung tumor burden by grade (lung tumor area/ total lung area per grade). *p<0.05 (unpaired t test with Holm-Sidak’s multiple comparisons test). For both **I** and **J** n=10 mice and ≥2,000 tumors per genotype.

### Nrf2 activation impairs lung adenocarcinoma progression

The decrease in grade 3 tumor burden in the Keap1/Nrf2 mutant models suggested that Nrf2 activation may impair progression to higher grade tumors. We next used the Kras^G12D/+^; p53^fl/fl^ (KP) model, which develops advanced-grade lung adenocarcinomas (43). We previously reported that the expression of Keap1^R554Q/R554Q^ dramatically decreased overall tumor size in this model (32), but other groups have found differing effects of Nrf2 activation. While some studies reported that Keap1 inactivation promoted adenocarcinoma progression (33, 34), others reported that Keap1 deletion did not affect tumor size(36, 37), but the conditions used and phenotypes assayed varied across these studies. Thus, we decided to perform comprehensive phenotyping on KP tumors following Nrf2 activation. Similar to what we observed in the Kras^G12D/+^ model, we found that overall survival of the KP model was not affected by Keap1 or Nrf2 mutation **(Fig. 5A** and **B)**. Moreover, we found that Keap1/Nrf2 mutation affected Nrf2 activation in a similar manner to the Kras^G12D/+^ model, with Keap1^R554Q/R554Q^ being the most activating, followed by Nrf2^D29H/+^, and then Keap1^R554Q/+^ compared to Keap1/Nrf2^+/+^ mice **(Fig. 5C-F)**. From the histology images, it was very apparent that Keap1^R554Q/R554Q^ mice had decreased overall tumor burden **(Fig. 5G)**, which was confirmed upon quantitation (Supplementary Fig. S3A). Next, we examined tumor progression by tumor grading and found a significant decrease in the proportion, number, and burden of grade 3 and 4 tumors in the Keap1^R554Q/R554Q^ cohort **(Fig. 5G** and **H**, Supplementary Fig. S3B). We also observed a modest, non-significant decrease in the proportion, number, and burden of grade 3 and 4 tumors in the Nrf2^D29H/+^ cohort **(Fig. 5G** and **H**, Supplementary Fig. S3B), further supporting a threshold for Nrf2 to impair tumor progression. Additionally, we saw an increase in the proportion, number, and burden of grade 1 tumors in the Keap1^R554Q/R554Q^ group, suggesting that there may be a threshold for Nrf2 activation to promote early progression, but impair late progression **(Fig. 5G** and **H**, Supplementary Fig. S3B). While there were no differences in overall median tumor size (Supplementary Fig. S3C), Keap1^R554Q/R554Q^ alveolar hyperplasia and grade 1 tumors were significantly smaller compared to their wild-type counterparts (Supplementary Fig. S3D). Consistently, we found a decrease in tumor cell proliferation in the Keap1^R554Q/R554Q^ cohort, specifically in AAH and grade 1/2 tumors, and no tumor cell death was observed (Supplementary Fig. S3E-H). Importantly, despite the reduction in grade 1 size, the higher grade 1 burden in Keap1^R554Q/R554Q^ mice was accounted for by the increase in the total number of grade 1 tumors (Supplementary Fig. S3B and I), highlighting that burden is a complex measurement influenced by multiple variables. We next investigated whether tumors that progressed to the adenocarcinoma stage altered Nrf2 expression and/or activity. To this end, we analyzed Nrf2 and Nqo1 levels across all tumor grades and hyperplasia (AAH, BH). We found that Nrf2 and Nqo1 levels were highly elevated in Keap1/Nrf2 mutant grade 1 tumors, with Nqo1 demonstrating increased nuclear localization in homozygous Keap1 mutant tumors compared to Nrf2 mutant tumors **(Fig. 6A)**. However, as tumors progressed to higher grades, Nrf2 and Nqo1 expression were reduced in the Keap1/Nrf2 mutant models **(Fig. 6A-C)**. These results suggest that Nrf2 activation beyond a certain threshold impairs advanced-grade tumor progression, requiring selection for a more tolerable level of Nrf2 expression and activity in high-grade tumors.

**Figure 5.**
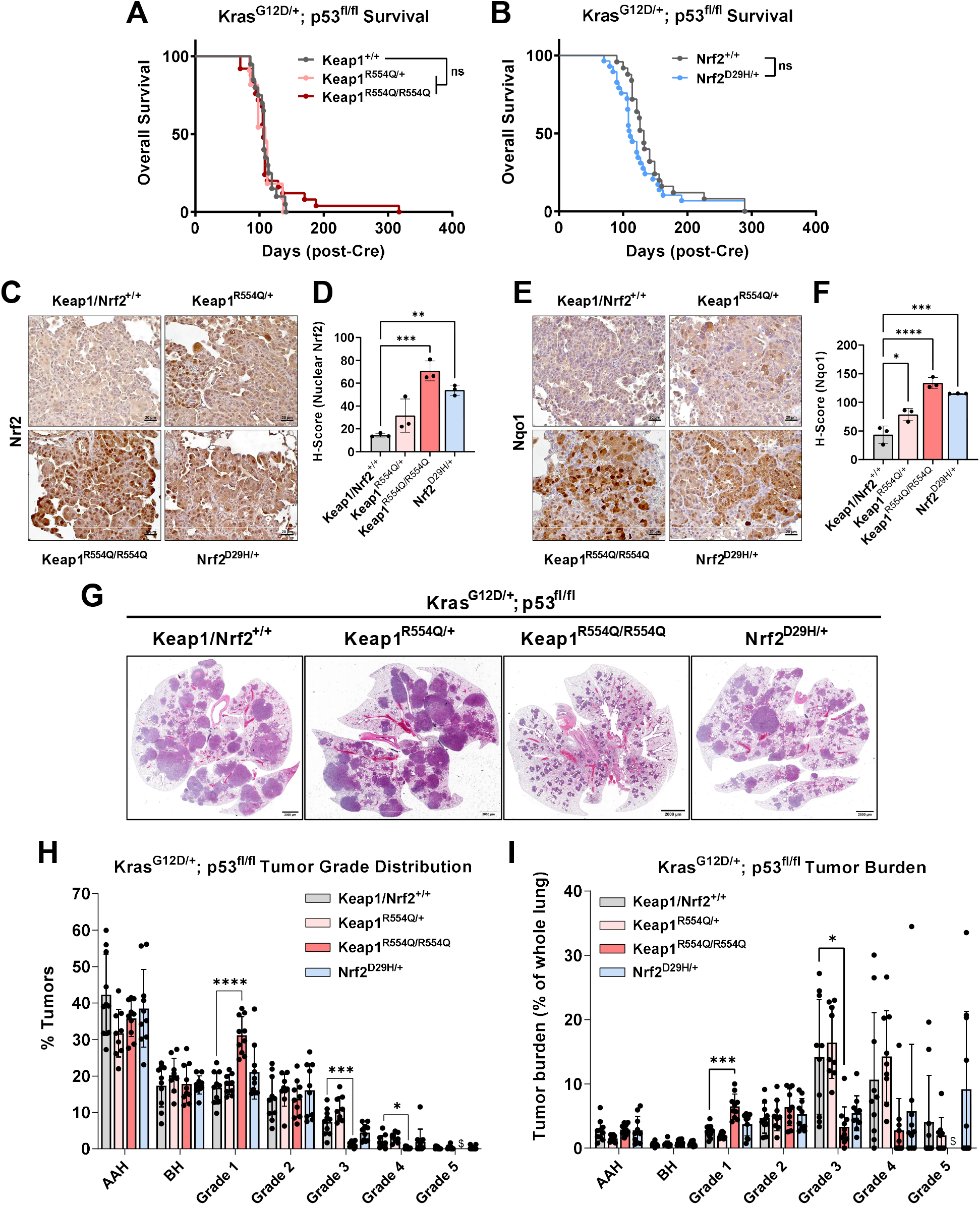
Homozygous Keap1^R554Q^ impairs adenocarcinoma progression in the Kras^G12D/+^; p53^fl/fl^ model. **A**, Overall survival of Kras^G12D/+^; p53^fl/fl^ mice with Keap1 mutation. Keap1^+/+^ (n=20); Keap1^R554Q/+^ (n=11); Keap1^R554Q/R554Q^ (n=25). **B**, Overall survival of Kras^G12D/+^; p53^fl/fl^ mice with Nrf2 mutation. Nrf2^+/+^ (n=25); Nrf2^D29H/+^ (n=29). Ns = not significant (Log-rank (Mantel-Cox) test). **C**, Representative immunohistochemical (IHC) staining of Nrf2 in Kras^G12D/+^; p53^fl/fl^ mice with Keap1/Nrf2 mutation (scale bars = 20 μM). **D**, H-scores for Nrf2 (nuclear) IHC staining. **E**, Representative immunohistochemical (IHC) staining of Nrf2 target Nqo1 (scale bars = 20 μM). **F**, H-scores for Nqo1 (whole cell) IHC staining. For **C-F**, N=3 mice per genotype and >20,000 tumor cells per mouse. *p<0.05 (one-way ANOVA). **G**, Representative whole lung H&E stained section (scale bars = 2000 μM). **H**, Distribution of tumor grades across Keap1/Nrf2 mutant models. *p<0.05 (unpaired t test with Holm-Sidak’s multiple comparisons test). $ = fewer than 3 tumors detected across all mice. **I**, Fraction of lung tumor burden by grade (lung tumor area per grade/ total lung area). *p<0.05 (unpaired t test with Holm-Sidak’s multiple comparisons test). $ = fewer than 3 tumors detected across all mice. For both **H** and **I** n≥9 mice and ≥1,900 tumors per genotype. Only one grade 5 tumor was found in the Keap1^R554Q/R554Q^ cohort, and therefore was excluded from these analyses.

**Figure 6.**
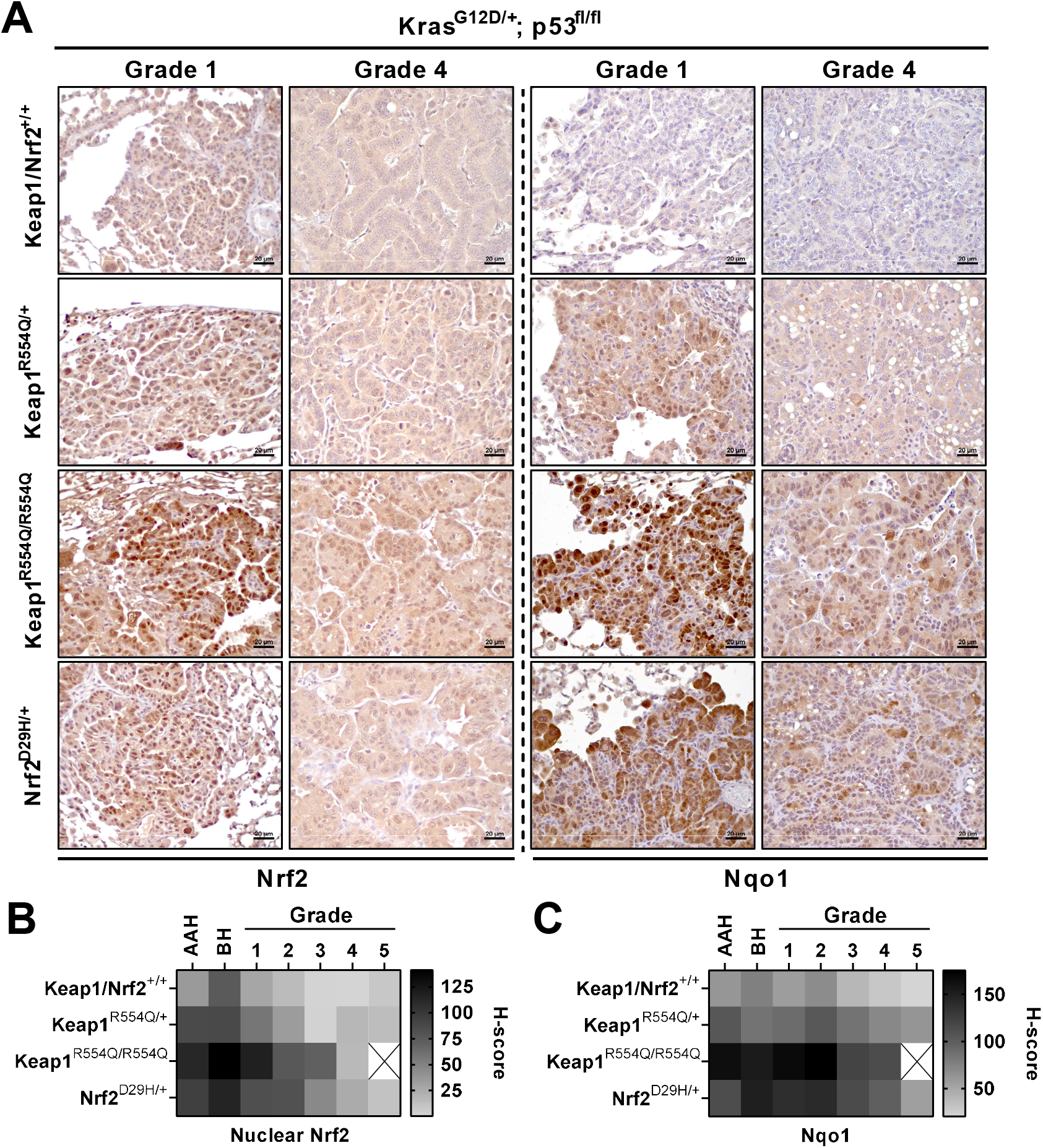
Nrf2 expression and activity is reduced in higher-grade tumors. **A**, Representative Nrf2 and Nqo1 IHC staining in grade 1 and 4 tumors from Kras^G12D/+^; p53^fl/fl^ mice with Keap1 or Nrf2 mutation (scale bars = 20 μM). **B, C** Heatmaps depicting the H-scores per grade from IHC staining for Nrf2 (nuclear) **(B)**, and the Nrf2 target Nqo1 (whole cell) **(C)**. N=3 mice per genotype, >20,000 tumor cells per mouse. Only one grade 5 tumor was found in the Keap1^R554Q/R554Q^ cohort, and therefore was excluded from these analyses.

### NRF2 overexpression impairs NSCLC cell proliferation, viability, and anchorage-independent colony formation

KEAP1 has other substrates (51–56), raising the question of whether NRF2 plays a causal role in tumor suppression. It was previously reported that KEAP1 mutant lung cancer cell lines are “NRF2 addicted” and dependent on NRF2 for proliferation (57). Supportingly, analysis of DepMap data (45) revealed that NSCLC cell lines with high NRF2 activity (24), which were enriched for KEAP1 mutations, exhibited NRF2 dependence **(Fig. 7A)**. To directly test the hypothesis that excessive NRF2 activation above a specific threshold is detrimental to lung tumor cell growth, we used lentiviral transduction to overexpress NRF2 in five KEAP1 mutant lung cancer cell lines (H1944, H322, A549, HCC15, H460). We confirmed overexpression of NRF2 by western blot analysis of NRF2 and target genes GCLC, xCT, and GSR **(Fig. 7B)**. Lentiviral transduction increased the expression of NRF2 in all cell lines, and increased the expression of NRF2 targets, demonstrating that NRF2 binding sites were not saturated by the level of NRF2 in these cell lines **(Fig. 7B)**. Next, we determined the influence of NRF2 overexpression on cellular proliferation and death over the course of four days using live cell imaging **(Fig. 7C-G)**. Like what we observed in the Kras^G12D/+^; p53^fl/fl^ model upon Keap1^R554Q/R554Q^ expression, we found that NRF2 overexpression decreased cell proliferation in all cell lines **(Fig. 7F)**. Moreover, Nrf2 overexpression increased cell death in 4 out of 5 cell lines **(Fig. 7G)**. Finally, we observed impaired anchorage-independent growth in soft agar in all cell lines **(Fig. 7H** and **I)**. These results indicate that there is an optimal threshold of NRF2 activity, and that excess NRF2 activation can impair lung cancer phenotypes.

**Figure 7.**
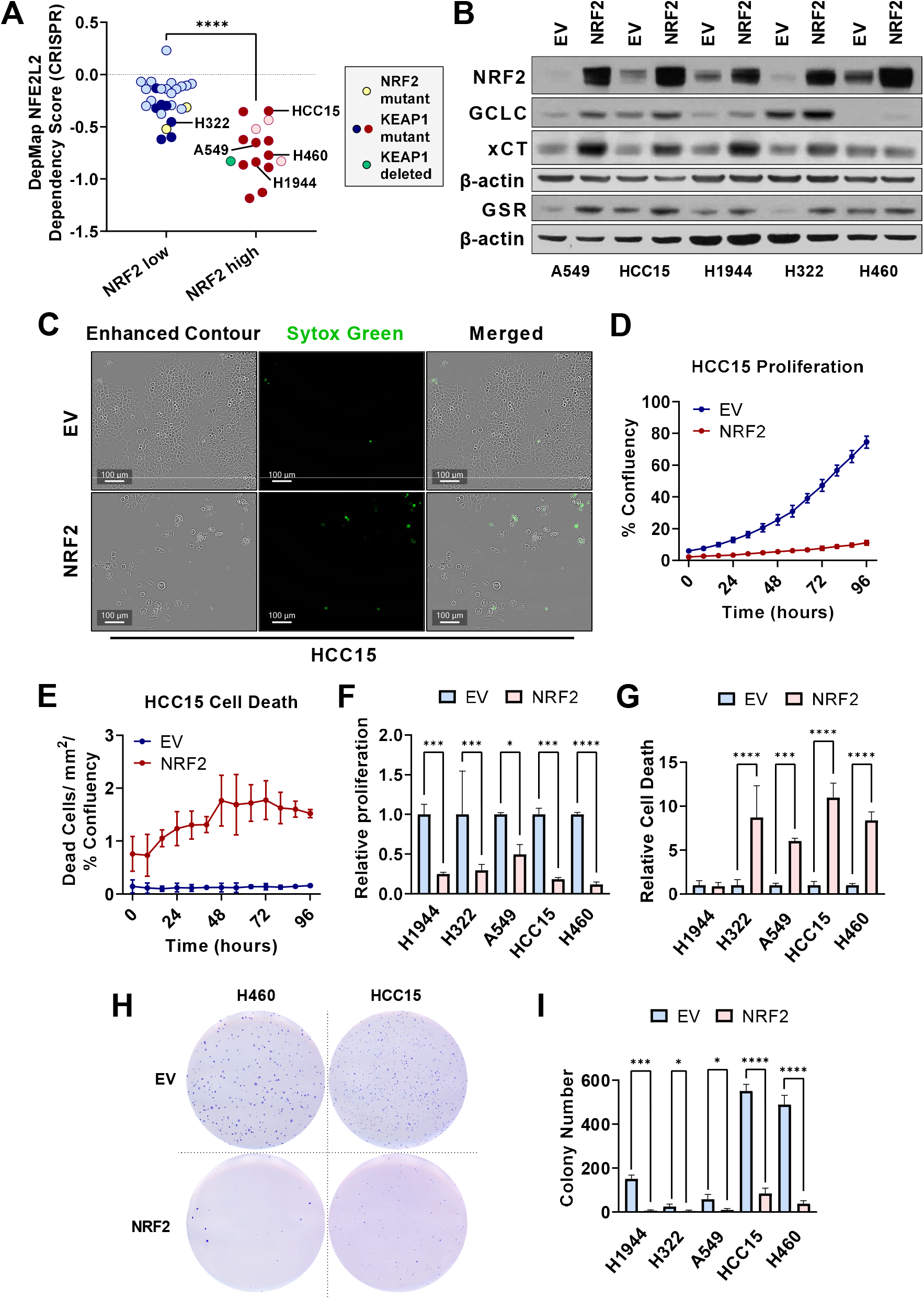
NRF2 overexpression impairs lung cancer cell proliferation, viability, and soft agar colony formation. **A**, Dependency scores obtained from DepMap(45) and represented as *NFEL2* 22Q2 Public+Score, Chronos for NSCLC cell lines previously determined to have high or low NRF2 activity(24). NRF2 mutant line symbols are represented by yellow, KEAP1 mutant lines by dark red or dark blue, and KEAP1 deleted lines by green. ***p<0.0001 (unpaired t-test). **B**, Western blot analysis of NRF2, β-actin, and NRF2 target GCLC, xCT, and GSR expression in *KEAP1* mutant lung cancer cell lines transduced with PLX317-empty vector (EV) or PLX317-NRF2 (NRF2). **C**, Representative images of HCC15 cells transduced with EV or NRF2 demonstrating cell confluency (enhanced contour) and cell death (Sytox Green) (scale bars = 100 μM). **D, E**, Analysis of EV and NRF2 HCC15 cell proliferation and death over 96 hours. Proliferation is represented as % confluency at each time point, and cell death as the number of Sytox Green positive cells per area normalized to % confluency. N=3 technical replicates per cell line, and two independent experiments. **F, G**, Area under the curve (AUC) analysis of cell proliferation **(F)** and Sytox Green-positive cell death **(G)** in KEAP1 mutant lung cancer cells lines +/− NRF2, normalized to empty vector control. *p<0.05 (one-way ANOVA). For **C-G** NSCLC cells were seeded in triplicate in 96-well plates at a density of 2,500 cells/ well. **H**, Representative images of H460 and HCC15 soft agar colony formation +/− NRF2. **I**, Quantification of soft agar colony number of *KEAP1* mutant lung cancer cell lines. *p<0.05 (one-way ANOVA). N=3 technical replicates per cell line, two independent experiments. For **H, I**, 5,000 cells per well were seeded in 6-well plates in triplicate.

### Single copy Nrf2 deletion rescues homozygous Keap1^R554Q^-mediated tumor progression impairment

To directly examine whether reducing Nrf2 levels could alleviate the block in adenocarcinoma progression in the Keap1^R554Q/R554Q^ model, we crossed a Nrf2^flox^ allele into both our Kras^G12D/+^; p53^fl/fl^ and Kras^G12D/+^; p53^fl/fl^; Keap1^R554Q/R554Q^ models. Because we previously found that complete Nrf2 deficiency impairs tumor initiation (23), we examined the consequence of single copy Nrf2 deletion on tumor phenotypes. Again, we found no difference in overall survival between groups **(Fig. 8A)**. We also observed that Nrf2 deletion in the Keap1^R554Q/R554Q^ model significantly decreased expression of Nrf2 and Nqo1 **(Fig. 8B-E**, Supplementary Fig. S4). Moreover, histological examination of the lungs revealed a striking difference in tumor burden, with Nrf2 heterozygous deletion having a minimal effect on the Kras^G12D/+^; p53^fl/fl^ model while dramatically increasing burden in the Kras^G12D/+^; p53^fl/fl^; Keap1^R554Q/R554Q^ model **(Fig. 8F**, Supplementary Fig. S5A). Tumor size was not affected (Supplementary Fig. S5B and C). Next, we examined tumor progression in these models. In agreement with our previous experiment **(Fig. 5H)**, we found that the Keap1^R554Q/R554Q^ cohort had a significant reduction in adenocarcinoma progression upon Nrf2^WT^ expression **(Fig. 8G)**. However, upon single copy Nrf2 deletion (Nrf2^flox/+^), Keap1^R554Q/R554Q^ failed to suppress tumor progression **(Fig. 8G)**. Moreover, the decrease in grade 3 tumor number and burden induced by Keap1^R554Q/R554Q^ was alleviated by single copy deletion of Nrf2 **(Fig. 8H**, Supplementary Fig. **S5D** and **E)**. These results demonstrate that there is a threshold by which Nrf2 activation can promote or impair tumor initiation or progression.

**Figure 8.**
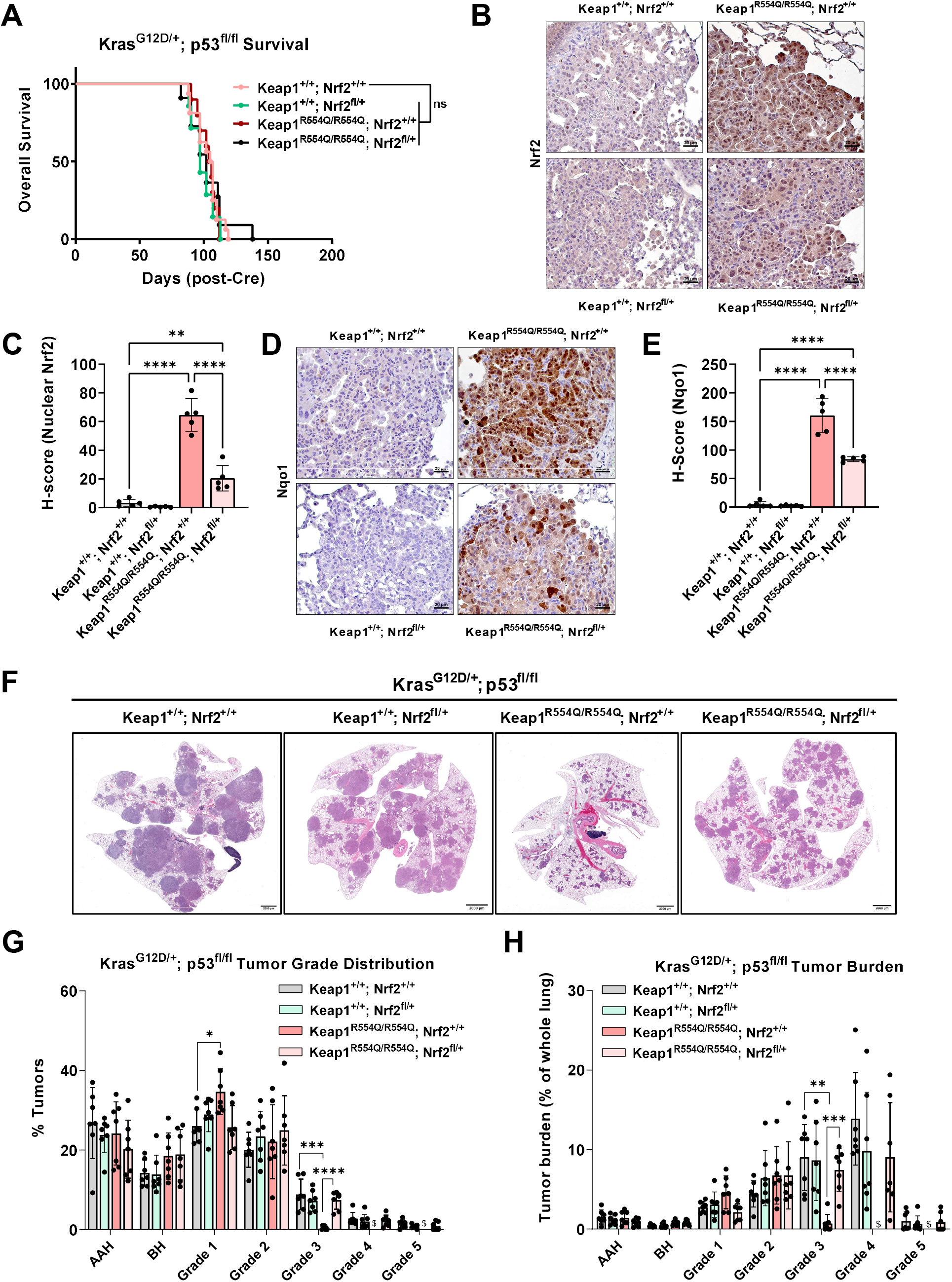
Single copy Nrf2 deletion rescues homozygous Keap1^R554Q^-mediated adenocarcinoma progression impairment in the Kras^G12D/+^; p53^fl/fl^ model. **A**, Overall survival of Kras^G12D/+^; p53^fl/fl^ mice with Keap1 mutation and/ or single copy Nrf2 deletion. Keap1^+/+^; Nrf2^+/+^ (n=16), Keap1^+/+^; Nrf2^fl/+^ (n=7), Keap1^R554Q/R554Q^; Nrf2^+/+^ (n=10), Keap1^R554Q/R554Q^; Nrf2^fl/+^ (n=11). Ns = not significant (Log-rank (Mantel-Cox) test). **B**, Representative immunohistochemical (IHC) staining of Nrf2 in Kras^G12D/+^; p53^fl/fl^ mice with Keap1 mutation and/ or heterozygous Nrf2 deletion (scale bars = 20 μM). **C**, H-scores for Nrf2 (nuclear) IHC staining. **D**, Representative IHC staining of Nrf2 target Nqo1 (scale bars = 20 μM). **E**, H-scores for Nqo1 (whole-cell) IHC staining. For **B-E**, N=3 mice per genotype and >20,000 tumor cells per mouse. *p<0.05 (one-way ANOVA). **F**, Representative whole lung H&E-stained sections (scale bars = 2000 μM). **G**, Distribution of tumor grades across Keap1 mutant/ Nrf2-deleted models. *p<0.05 (unpaired t test with Holm-Sidak’s multiple comparisons test). $ = fewer than 3 tumors detected across all mice. **H**, Fraction of lung tumor burden by grade (lung tumor area per grade/ total lung area). *p<0.05 (unpaired t test with Holm-Sidak’s multiple comparisons test). $ = fewer than 3 tumors detected across all mice. For both **G** and **H** n=7 mice and ≥1,000 tumors per genotype were analyzed. Only one grade 4 and one grade 5 tumor were found in the Keap1^R554Q/R554Q^ cohort, and therefore were excluded from these analyses.

## Discussion

Using genetically engineered mouse models of Keap1/Nrf2 mutation, we find that Keap1 or Nrf2 mutations alone are insufficient to cause lung tumor formation. Even in combination with tumor suppressor loss, we did not observe lung tumor formation after 500 days, suggesting that chronic NRF2 activation would be a safe strategy for chemoprevention. These findings corroborate previous studies showing that Keap1 deletion does not induce lung tumor development (28), even in the absence of tumor suppressors p53 or Lkb1 (29), out to 12-15 months. In combination with Kras^G12D/+^ mutation, we found that Nrf2 activation promoted tumor initiation. Our results are consistent with previous work indicating that Nrf2 activation via Keap1 deletion promotes Kras^G12D/+^ tumor initiation (29, 30). Surprisingly, we find that Nrf2 activation impairs tumor progression, which is correlated with Nrf2 dosage. In our previous study with Keap1^R554Q/R554Q^ mice, we reported smaller tumors in the KP model (32), which we now find is due to impaired tumor progression mediated by Nrf2 hyperactivation. Because most prior studies from other groups did not specifically analyze tumor grade (27, 29, 36, 37), they may not have captured this effect of Nrf2 on tumor progression. Alternatively, there may be biological differences between Keap1 deletion and mutation. Moreover, the dosage-dependent effects of Nrf2 that we found are also reminiscent of what has been observed for oncogenes like Ras and Myc, where low levels promote transformation and proliferation, and high levels promote senescence or death (58, 59).

Our findings that tumors downregulate Nrf2 to select for a level permissive for tumor progression and that ectopic NRF2 expression antagonizes the proliferation and viability of human NSCLC cells are supported by our finding that single copy deletion of Nrf2 can rescue the homozygous Keap1^R554Q^-mediated block in tumor progression. This result demonstrates a direct role for Nrf2 hyperactivation but we cannot exclude the possibility that alternative KEAP1 substrates, such as PGAM5 (51), PALB2 (52), MCM3 (53) or EMSY (54), contribute to the block in tumor progression. In agreement with our findings, a Keap1-binding defective Nrf2^E79Q^ mouse model of small-cell lung cancer (SCLC) with p53/ p16 inactivation (60) also displayed Nrf2 downregulation in aggressive SCLC tumors. The exact mechanism(s) by which high NRF2 activity impairs tumor cell proliferation and tumor progression remains to be determined. NRF2 activation is associated with multiple metabolic liabilities that could explain this phenotype, including CDO1-mediated toxic metabolite formation and NADPH depletion (32), xCT-mediated glutamate depletion (26, 34, 61), and ALDH3A1-mediated reductive stress (62). Future studies are needed to determine if these or other mechanisms play a role in NRF2-mediated proliferation and progression impairment.

Given that KEAP1 and NRF2 mutations are found with a high frequency in human NSCLC and are associated with poor outcomes, these findings raise the question of under which contexts NRF2 activation provides an advantage. Our findings that NRF2/KEAP1 mutation promotes tumor initiation are consistent with recent results from the TRACERx study, where *KEAP1* mutations were found to be an initiating driver together with *KRAS* and *TP53* mutations in lung adenocarcinoma (63), and suggest that additional genetic events may be needed to overcome NRF2-mediated inhibition of tumor progression. We did not find that these mutations conferred poor outcomes in our mouse models, but there are multiple features of patient tumors not captured by our models that remain to be examined. They did not develop metastases with sufficient frequency, precluding an examination of the influence of Nrf2 activation on metastasis as reported previously (27). Moreover, mice are not exposed to smoking and other environmental toxins under which NRF2 activation may promote survival (4–7). We have also not tested the response of these models to therapy. Additional mechanistic studies are warranted to clarify the clinical translation of these approaches, including understanding how NRF2 is downregulated upon progression and the consequences of NRF2 pharmacological activation and inhibition on the various tumor grades in our GEMM models. These studies will be important to compare the level of NRF2 achieved with pharmacological manipulation to the level induced by KEAP1/NRF2 mutation and where that falls on the threshold spectrum. Moreover, our models are designed to only activate NRF2 in cancer cells while these pharmacological agents will modulate NRF2 in the microenvironment, which will also influence tumor growth (30, 64, 65). These future studies will further clarify the context-dependent role of NRF2 during the complex stages of tumorigenesis.

## Acknowledgements

This work was supported by grants from the NIH/NCI (R37-CA230042), the American Lung Association (LCDA-498544) and the American Cancer Society’s Institutional Research Grant to G.M.D. This work was also supported by the Analytical Microscopy and the Tissue Core Facilities at the Moffitt Cancer Center, an NCI designated Comprehensive Cancer Center (P30-CA076292). We would like to thank the DeNicola lab members and Dr. Ana Gomes for the helpful discussions. We thank Cheyenne Schneider for proofreading.

## Online supplemental material

Supplementary Fig. S1 shows that the *CA-Keap1*^*R554Q*^ allele does not exhibit hypomorphism in mouse embryonic fibroblasts. Supplementary Fig. S2 shows lung tumor analysis data for Keap1/Nrf2 mutant mice in the Kras^G12D/+^ model. Supplementary Fig. S3 shows lung tumor analysis data and tumor cell proliferation/ death immunohistochemical staining for Keap1/Nrf2 mutant mice in the Kras^G12D/+^; p53^fl/fl^ model. Supplementary Fig. S4 shows the H-scores by tumor grade of Nrf2 and Nqo1 immunohistochemical staining in Kras^G12D/+^; p53^fl/fl^ and Kras^G12D/+^; p53^fl/fl^; Keap1^R554Q/R554Q^ mouse models with single copy Nrf2 deletion. Supplementary Fig. S5 shows lung tumor analysis data in the Kras^G12D/+^; p53^fl/fl^ model with single copy Nrf2 deletion.

## Author Contributions

G.M.D. conceived the project and G.M.D. and J.M.D. designed the experiments. G.M.D., I.I.C.C. and F.A.K. designed and generated the Nrf2 and Keap1 mutant mice. G.M.D. performed the RNA sequencing. Maintenance of animal colonies, generation of experimental animals, and collections of tissues were performed by A.F., with assistance from S.C. and N.F.P. J.M.D. performed cell line experiments, immunohistochemistry, and tumor histology analyses. J.R.P. and E.E.S. generated the affinity-purified NRF2 antibody used for immunohistochemistry. J.M.D. and G.M.D. wrote the manuscript, and all authors reviewed it. G.M.D. acquired funding and supervised the study.

## References

1. DeBlasi JM, DeNicola GM. Dissecting the Crosstalk between NRF2 Signaling and Metabolic Processes in Cancer. Cancers (Basel). 2020;12(10).

2. He F, Ru X, Wen T. NRF2, a Transcription Factor for Stress Response and Beyond. Int J Mol Sci. 2020;21(13).

3. Itoh K, Wakabayashi N, Katoh Y, Ishii T, Igarashi K, Engel JD, et al. Keap1 represses nuclear activation of antioxidant responsive elements by Nrf2 through binding to the amino-terminal Neh2 domain. Genes Dev. 1999;13(1):76–86.

4. Ramos-Gomez M, Dolan PM, Itoh K, Yamamoto M, Kensler TW. Interactive effects of nrf2 genotype and oltipraz on benzo[a]pyrene–DNA adducts and tumor yield in mice. Carcinogenesis. 2003;24(3):461–7.

5. Ramos-Gomez M, Kwak M-K, Dolan PM, Itoh K, Yamamoto M, Talalay P, et al. Sensitivity to carcinogenesis is increased and chemoprotective efficacy of enzyme inducers is lost in <i>nrf2</i> transcription factor-deficient mice. Proceedings of the National Academy of Sciences. 2001;98(6):3410–5.

6. Aoki Y, Sato H, Nishimura N, Takahashi S, Itoh K, Yamamoto M. Accelerated DNA adduct formation in the lung of the Nrf2 knockout mouse exposed to diesel exhaust. Toxicol Appl Pharmacol. 2001;173(3):154–60.

7. Iida K, Itoh K, Kumagai Y, Oyasu R, Hattori K, Kawai K, et al. Nrf2 Is Essential for the Chemopreventive Efficacy of Oltipraz against Urinary Bladder Carcinogenesis. Cancer Research. 2004;64(18):6424–31.

8. Kensler TW, Wakabayashi N. Nrf2: friend or foe for chemoprevention? Carcinogenesis. 2010;31(1):90–9.

9. Schmidlin CJ, Shakya A, Dodson M, Chapman E, Zhang DD. The intricacies of NRF2 regulation in cancer. Semin Cancer Biol. 2021;76:110–9.

10. Telkoparan-Akillilar P, Suzen S, Saso L. Pharmacological Applications of Nrf2 Inhibitors as Potential Antineoplastic Drugs. Int J Mol Sci. 2019;20(8).

11. Network CGAR. Comprehensive molecular profiling of lung adenocarcinoma. Nature. 2014;511(7511):543–50.

12. Network CGAR. Comprehensive genomic characterization of squamous cell lung cancers. Nature. 2012;489(7417):519–25.

13. Singh A, Misra V, Thimmulappa RK, Lee H, Ames S, Hoque MO, et al. Dysfunctional KEAP1-NRF2 interaction in non-small-cell lung cancer. PLoS Med. 2006;3(10):e420.

14. Shibata T, Ohta T, Tong KI, Kokubu A, Odogawa R, Tsuta K, et al. Cancer related mutations in NRF2 impair its recognition by Keap1-Cul3 E3 ligase and promote malignancy. Proc Natl Acad Sci U S A. 2008;105(36):13568–73.

15. Wang R, An J, Ji F, Jiao H, Sun H, Zhou D. Hypermethylation of the Keap1 gene in human lung cancer cell lines and lung cancer tissues. Biochem Biophys Res Commun. 2008;373(1):151–4.

16. Kim YR, Oh JE, Kim MS, Kang MR, Park SW, Han JY, et al. Oncogenic NRF2 mutations in squamous cell carcinomas of oesophagus and skin. J Pathol. 2010;220(4):446–51.

17. Zhang P, Singh A, Yegnasubramanian S, Esopi D, Kombairaju P, Bodas M, et al. Loss of Kelch-like ECH-associated protein 1 function in prostate cancer cells causes chemoresistance and radioresistance and promotes tumor growth. Mol Cancer Ther. 2010;9(2):336–46.

18. Solis LM, Behrens C, Dong W, Suraokar M, Ozburn NC, Moran CA, et al. Nrf2 and Keap1 abnormalities in non-small cell lung carcinoma and association with clinicopathologic features. Clin Cancer Res. 2010;16(14):3743–53.

19. Wang Q, Xu L, Wang G, Chen L, Li C, Jiang X, et al. Prognostic and clinicopathological significance of NRF2 expression in non-small cell lung cancer: A meta-analysis. PLoS One. 2020;15(11):e0241241.

20. Zhao J, Lin X, Meng D, Zeng L, Zhuang R, Huang S, et al. Nrf2 Mediates Metabolic Reprogramming in Non-Small Cell Lung Cancer. Front Oncol. 2020;10:578315.

21. Sporn MB, Liby KT. NRF2 and cancer: the good, the bad and the importance of context. Nat Rev Cancer. 2012;12(8):564–71.

22. Singh A, Bodas M, Wakabayashi N, Bunz F, Biswal S. Gain of Nrf2 function in non-small-cell lung cancer cells confers radioresistance. Antioxidants and Redox Signaling. 2010.

23. Denicola GM, Karreth FA, Humpton TJ, Gopinathan A, Wei C, Frese K, et al. Oncogene-induced Nrf2 transcription promotes ROS detoxification and tumorigenesis. Nature. 2011;475(7354):106–10.

24. DeNicola GM, Chen PH, Mullarky E, Sudderth JA, Hu Z, Wu D, et al. NRF2 regulates serine biosynthesis in non-small cell lung cancer. Nature Genetics. 2015;47(12):1475–81.

25. Mitsuishi Y, Taguchi K, Kawatani Y, Shibata T, Nukiwa T, Aburatani H, et al. Nrf2 redirects glucose and glutamine into anabolic pathways in metabolic reprogramming. Cancer Cell. 2012;22(1):66–79.

26. Sayin VI, LeBoeuf SE, Singh SX, Davidson SM, Biancur D, Guzelhan BS, et al. Activation of the NRF2 antioxidant program generates an imbalance in central carbon metabolism in cancer. Elife. 2017;6.

27. Lignitto L, LeBoeuf SE, Homer H, Jiang S, Askenazi M, Karakousi TR, et al. Nrf2 Activation Promotes Lung Cancer Metastasis by Inhibiting the Degradation of Bach1. Cell. 2019;178(2):316-29.e18.

28. Best SA, De Souza DP, Kersbergen A, Policheni AN, Dayalan S, Tull D, et al. Synergy between the KEAP1/NRF2 and PI3K Pathways Drives Non-Small-Cell Lung Cancer with an Altered Immune Microenvironment. Cell Metabolism. 2018;27(4):935-43.e4.

29. Best SA, Ding S, Kersbergen A, Dong X, Song JY, Xie Y, et al. Distinct initiating events underpin the immune and metabolic heterogeneity of KRAS-mutant lung adenocarcinoma. Nature Communications. 2019;10(1):1–14.

30. Hayashi M, Kuga A, Suzuki M, Panda H, Kitamura H, Motohashi H, et al. Microenvironmental Activation of Nrf2 Restricts the Progression of Nrf2-Activated Malignant Tumors. Cancer Res. 2020;80(16):3331–44.

31. Jeong Y, Hoang NT, Lovejoy A, Stehr H, Newman AM, Gentles AJ, et al. Role of KEAP1/NRF2 and TP53 mutations in lung squamous cell carcinoma development and radiation resistance. Cancer Discovery. 2017;7(1):86–101.

32. Kang YP, Torrente L, Falzone A, Elkins CM, Liu M, Asara JM, et al. Cysteine dioxygenase 1 is a metabolic liability for non-small cell lung cancer. eLife. 2019.

33. Romero R, Sánchez-Rivera FJ, Westcott PMK, Mercer KL, Bhutkar A, Muir A, et al. Keap1 mutation renders lung adenocarcinomas dependent on Slc33a1. Nature Cancer. 2020;1(6):589–602.

34. Romero R, Sayin VI, Davidson SM, Bauer MR, Singh SX, Leboeuf SE, et al. Keap1 loss promotes Kras-driven lung cancer and results in dependence on glutaminolysis. Nature Medicine. 2017.

35. Singh A, Daemen A, Nickles D, Jeon S-M, Foreman O, Sudini K, et al. NRF2 Activation Promotes Aggressive Lung Cancer and Associates with Poor Clinical Outcomes. Clinical Cancer Research. 2021;27(3):877.

36. Foggetti G, Li C, Cai H, Hellyer JA, Lin W-Y, Ayeni D, et al. Genetic Determinants of EGFR-Driven Lung Cancer Growth and Therapeutic Response In Vivo. Cancer Discovery. 2021;11(7):1736–53.

37. Rogers ZN, McFarland CD, Winters IP, Seoane JA, Brady JJ, Yoon S, et al. Mapping the in vivo fitness landscape of lung adenocarcinoma tumor suppression in mice. Nature Genetics. 2018.

38. Cai H, Chew SK, Li C, Tsai MK, Andrejka L, Murray CW, et al. A Functional Taxonomy of Tumor Suppression in Oncogenic KRAS-Driven Lung Cancer. Cancer Discov. 2021;11(7):1754–73.

39. Rogers ZN, McFarland CD, Winters IP, Naranjo S, Chuang C-H, Petrov D, et al. A quantitative and multiplexed approach to uncover the fitness landscape of tumor suppression in vivo. Nature Methods. 2017;14(7):737–42.

40. Jackson EL, Willis N, Mercer K, Bronson RT, Crowley D, Montoya R, et al. Analysis of lung tumor initiation and progression using conditional expression of oncogenic K-ras. Genes and Development. 2001.

41. McLoughlin MR, Orlicky DJ, Prigge JR, Krishna P, Talago EA, Cavigli IR, et al. TrxR1, Gsr, and oxidative stress determine hepatocellular carcinoma malignancy. Proc Natl Acad Sci U S A. 2019;116(23):11408–17.

42. Bankhead P, Loughrey MB, Fernandez JA, Dombrowski Y, McArt DG, Dunne PD, et al. QuPath: Open source software for digital pathology image analysis. Sci Rep. 2017;7(1):16878.

43. Jackson EL, Olive KP, Tuveson DA, Bronson R, Crowley D, Brown M, et al. The differential effects of mutant p53 alleles on advanced murine lung cancer. Cancer Research. 2005;65(22):10280–8.

44. Berger AH, Brooks AN, Wu X, Shrestha Y, Chouinard C, Piccioni F, et al. High-throughput Phenotyping of Lung Cancer Somatic Mutations. Cancer Cell. 2016;30(2):214–28.

45. Institute B. DepMap Public 22Q2 2022.

46. Hast BE, Cloer EW, Goldfarb D, Li H, Siesser PF, Yan F, et al. Cancer-derived mutations in KEAP1 impair NRF2 degradation but not ubiquitination. Cancer Research. 2014.

47. Tong KI, Padmanabhan B, Kobayashi A, Shang C, Hirotsu Y, Yokoyama S, et al. Different Electrostatic Potentials Define ETGE and DLG Motifs as Hinge and Latch in Oxidative Stress Response. Molecular and Cellular Biology. 2007.

48. Taguchi K, Maher JM, Suzuki T, Kawatani Y, Motohashi H, Yamamoto M. Genetic Analysis of Cytoprotective Functions Supported by Graded Expression of Keap1. Molecular and Cellular Biology. 2010.

49. Wakabayashi N, Itoh K, Wakabayashi J, Motohashi H, Noda S, Takahashi S, et al. Keap1-null mutation leads to postnatal lethality due to constitutive Nrf2 activation. Nature Genetics. 2003.

50. Huppke P, Weissbach S, Church JA, Schnur R, Krusen M, Dreha-Kulaczewski S, et al. Activating de novo mutations in NFE2L2 encoding NRF2 cause a multisystem disorder. Nature Communications. 2017;8(1):818.

51. Lo SC, Hannink M. PGAM5 tethers a ternary complex containing Keap1 and Nrf2 to mitochondria. Exp Cell Res. 2008;314(8):1789–803.

52. Ma J, Cai H, Wu T, Sobhian B, Huo Y, Alcivar A, et al. PALB2 interacts with KEAP1 to promote NRF2 nuclear accumulation and function. Mol Cell Biol. 2012;32(8):1506–17.

53. Mulvaney KM, Matson JP, Siesser PF, Tamir TY, Goldfarb D, Jacobs TM, et al. Identification and characterization of MCM3 as a kelch-like ECH-associated protein 1 (KEAP1) substrate. Journal of Biological Chemistry. 2016;291(45):23719–33.

54. Marzio A, Kurz E, Sahni JM, Di Feo G, Puccini J, Jiang S, et al. EMSY inhibits homologous recombination repair and the interferon response, promoting lung cancer immune evasion. Cell. 2022;185(1):169-83.e19.

55. Zhang Y, Shi Z, Zhou Y, Xiao Q, Wang H, Peng Y. Emerging Substrate Proteins of Kelch-like ECH Associated Protein 1 (Keap1) and Potential Challenges for the Development of Small-Molecule Inhibitors of the Keap1-Nuclear Factor Erythroid 2-Related Factor 2 (Nrf2) Protein– Protein Interaction. Journal of Medicinal Chemistry. 2020;1.

56. McCutcheon DC, Lee G, Carlos A, Montgomery JE, Moellering RE. Photoproximity Profiling of Protein-Protein Interactions in Cells. J Am Chem Soc. 2020;142(1):146–53.

57. Kitamura H, Motohashi H. NRF2 addiction in cancer cells. Cancer Sci. 2018;109(4):900–11.

58. Tuveson DA, Shaw AT, Willis NA, Silver DP, Jackson EL, Chang S, et al. Endogenous oncogenic K-ras(G12D) stimulates proliferation and widespread neoplastic and developmental defects. Cancer Cell. 2004;5(4):375–87.

59. Murphy DJ, Junttila MR, Pouyet L, Karnezis A, Shchors K, Bui DA, et al. Distinct thresholds govern Myc’s biological output in vivo. Cancer Cell. 2008;14(6):447–57.

60. Hamad SH, Montgomery SA, Simon JM, Bowman BM, Spainhower KB, Murphy RM, et al. TP53, CDKN2A/P16, and NFE2L2/NRF2 regulate the incidence of pure- and combined-small cell lung cancer in mice. Oncogene. 2022;41(25):3423–32.

61. Muir A, Danai LV, Gui DY, Waingarten CY, Lewis CA, Vander Heiden MG. Environmental cystine drives glutamine anaplerosis and sensitizes cancer cells to glutaminase inhibition. Elife. 2017;6.

62. Weiss-Sadan T, Ge M, Hayashi M, Gohar M, Yao C-H, de Groot A, et al. NRF2 activation induces NADH-reductive stress, providing a metabolic vulnerability in lung cancer. Cell Metabolism. 2023.

63. Jamal-Hanjani M, Wilson GA, McGranahan N, Birkbak NJ, Watkins TBK, Veeriah S, et al. Tracking the Evolution of Non–Small-Cell Lung Cancer. New England Journal of Medicine. 2017;376(22):2109–21.

64. Satoh H, Moriguchi T, Saigusa D, Baird L, Yu L, Rokutan H, et al. NRF2 Intensifies Host Defense Systems to Prevent Lung Carcinogenesis, but After Tumor Initiation Accelerates Malignant Cell Growth. Cancer Res. 2016;76(10):3088–96.

65. Satoh H, Moriguchi T, Taguchi K, Takai J, Maher JM, Suzuki T, et al. Nrf2-deficiency creates a responsive microenvironment for metastasis to the lung. Carcinogenesis. 2010;31(10):1833–43.

