## Supplemental Figures for "Distinct Nrf2 Signaling Thresholds Mediate Lung Tumor Initiation and Progression"

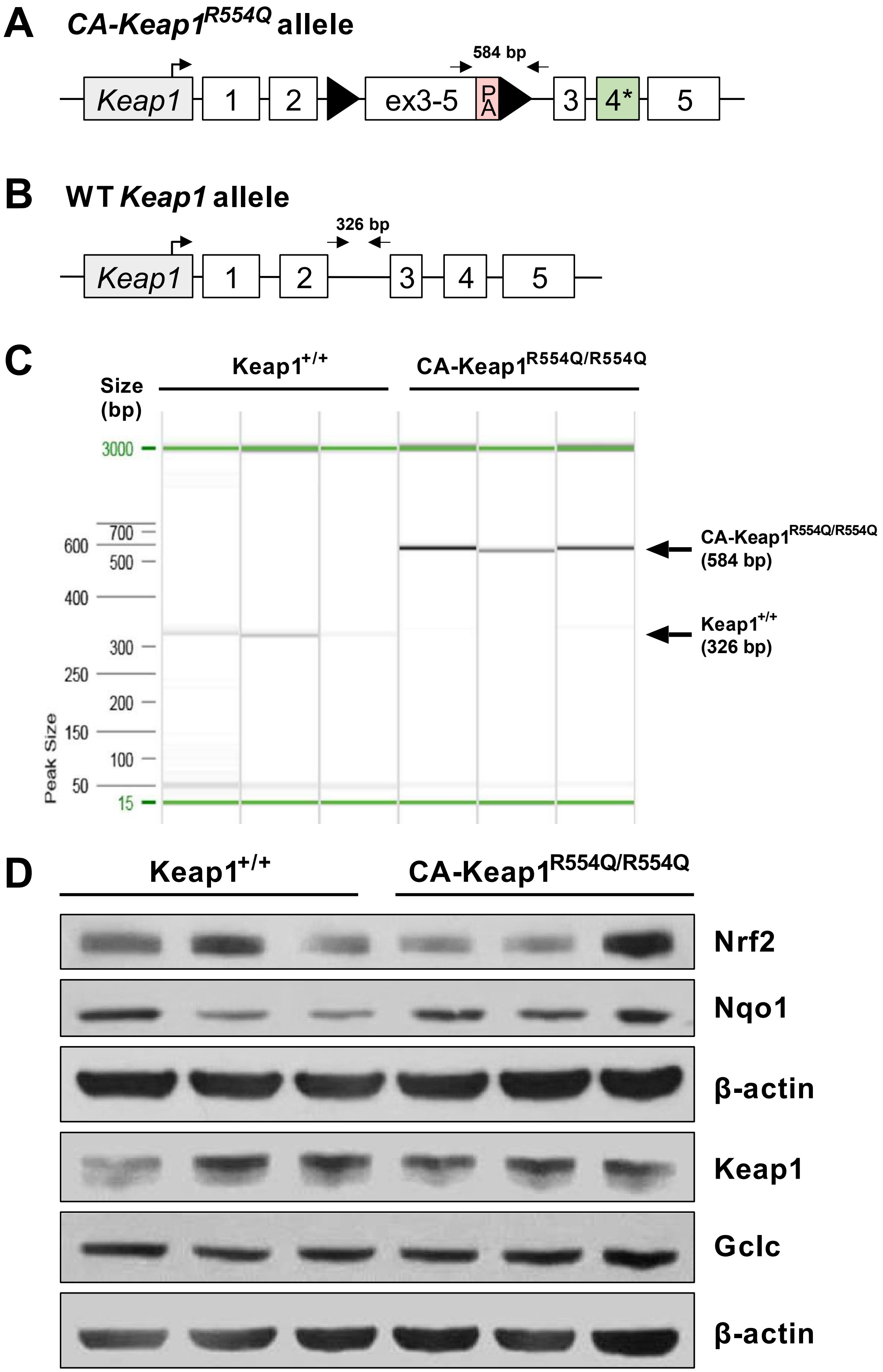
A
CA-Keap1R554Q allele
584 bp
Keap1
1
2
ex3-5
PA
3
4*
5
B
WT Keap1 allele
326 bp
Keap1
1
2
3
4
5
C
Keap1+/+
CA-Keap1R554Q/R554Q
Size (bp)
3000
700
600
500
400
300
250
200
150
100
50
15
Peak Size
CA-Keap1R554Q/R554Q (584 bp)
Keap1+/+ (326 bp)
D
Keap1+/+
CA-Keap1R554Q/R554Q
Nrf2
Nqo1
β-actin
Keap1
Gclc
β-actin

**Figure S1. The *CA-Keap1*$^{R554Q}$ allele is not hypomorphic.** Schematic of **(A)** *CA-Keap1*$^{R554Q}$ and **(B)** WT *Keap1* alleles with genotyping primers. **(C)** Detection of *CA-Keap1*$^{R554Q}$ and WT *Keap1* by QIAxcel electrophoresis. **D)** Western blot analysis of Nrf2, Nqo1, Keap1, Gclc, and β-actin expression in WT *Keap1* and *CA-Keap1*$^{R554Q/R554Q}$ MEFs (n=3 individual MEF lines from each genotype).

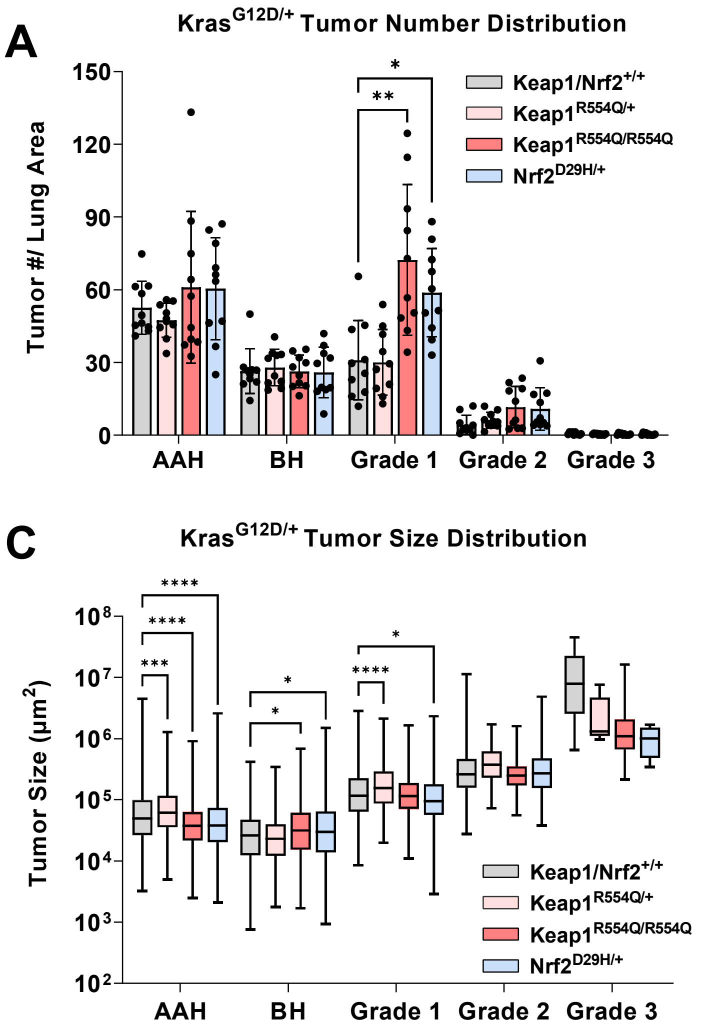

A
Kras$^{G12D/+}$ Tumor Number Distribution
Tumor #/ Lung Area
150
120
90
60
30
0
AAH
BH
Grade 1
Grade 2
Grade 3
*
**
Keap1/Nrf2$^{+/+}$
Keap1$^{R554Q/+}$
Keap1$^{R554Q/R554Q}$
Nrf2$^{D29H/+}$
B
Median Tumor Size (μm$^2$)
1.5×10$^5$
1×10$^5$
5×10$^4$
0
Keap1/Nrf2$^{+/+}$
Keap1$^{R554Q/+}$
Keap1$^{R554Q/R554Q}$
Nrf2$^{D29H/+}$
C
Kras$^{G12D/+}$ Tumor Size Distribution
Tumor Size (μm$^2$)
10$^8$
10$^7$
10$^6$
10$^5$
10$^4$
10$^3$
10$^2$
****
****
***
*
*
****
*
AAH
BH
Grade 1
Grade 2
Grade 3
Keap1/Nrf2$^{+/+}$
Keap1$^{R554Q/+}$
Keap1$^{R554Q/R554Q}$
Nrf2$^{D29H/+}$
D
Tumor Burden (%)
60
40
20
0
*
*
***
Keap1/Nrf2$^{+/+}$
Keap1$^{R554Q/+}$
Keap1$^{R554Q/R554Q}$
Nrf2$^{D29H/+}$

**Figure S2. Lung tumor burden, size, and number quantification in the Kras$^{G12D/+}$ model with Keap1/Nrf2 mutation. (A)** Distribution of tumor number by grade across Keap1/Nrf2 mutant models. N=10 mice per genotype, *p<0.05 (unpaired t test with Holm-Sidak's multiple comparisons test). **(B)** Median tumor size per mouse for each genotype. *p<0.05 (one-way ANOVA). **(C)** Tumor size by grade across all mice per genotype. N=10 mice and ≥2,300 tumors per genotype, *p<0.05 (Kruskal-Wallis test with Dunn's multiple comparisons test). **(D)** Overall tumor burden (%) calculated by dividing the total area of lung tumor by the total area of the lung. *p<0.05 (one-way ANOVA).

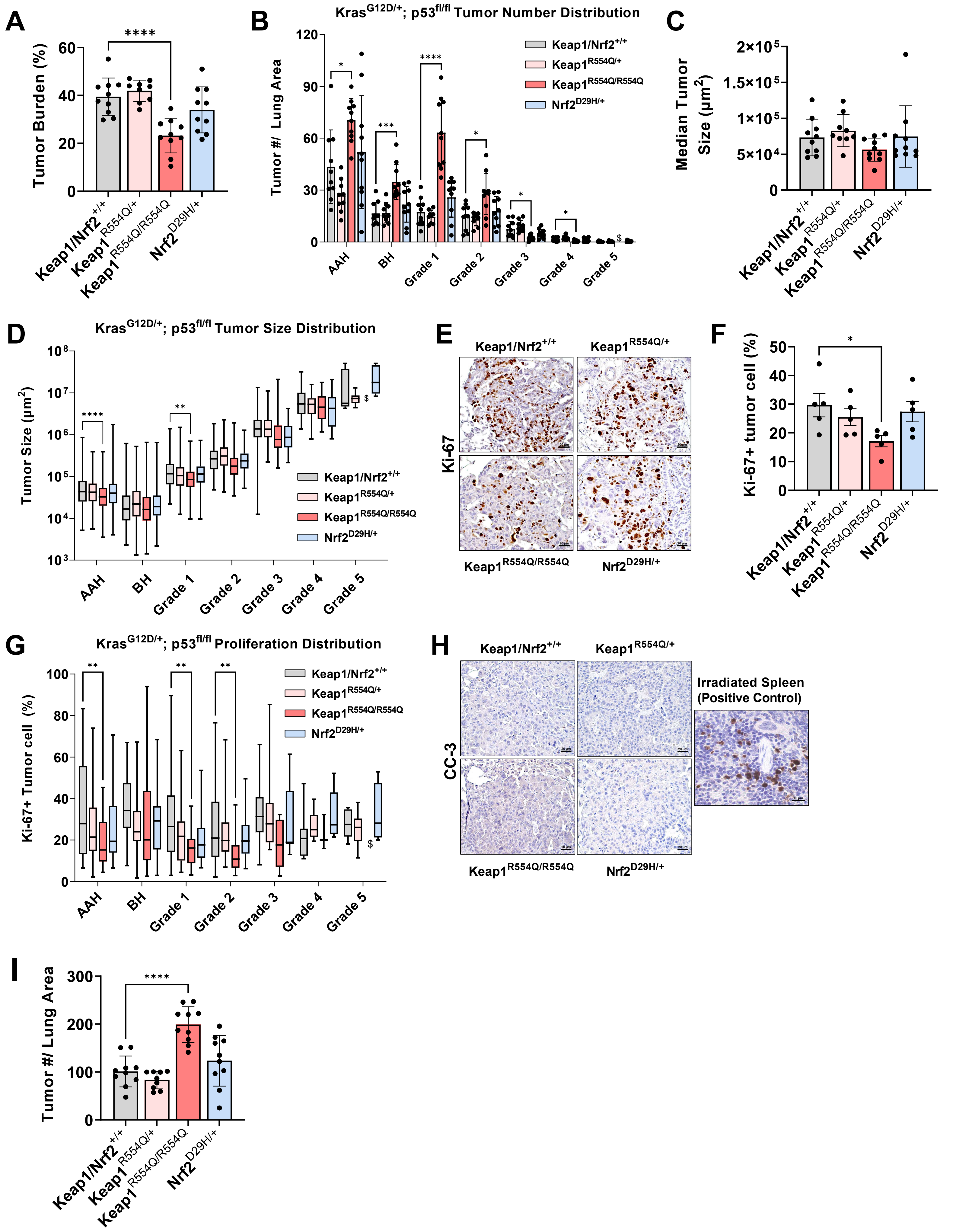
A
Tumor Burden (%)
****
Keap1/Nrf2+/+
Keap1R554Q/+
Keap1R554Q/R554Q
Nrf2D29H/+
B
KrasG12D/+; p53fl/fl Tumor Number Distribution
Tumor #/ Lung Area
Keap1/Nrf2+/+
Keap1R554Q/+
Keap1R554Q/R554Q
Nrf2D29H/+
AAH
BH
Grade 1
Grade 2
Grade 3
Grade 4
Grade 5
C
Median Tumor Size (μm2)
D
KrasG12D/+; p53fl/fl Tumor Size Distribution
Tumor Size (μm2)
E
Ki-67
F
Ki-67+ tumor cell (%)
G
KrasG12D/+; p53fl/fl Proliferation Distribution
Ki-67+ Tumor cell (%)
H
CC-3
Irradiated Spleen
(Positive Control)
I
Tumor #/ Lung Area

**Figure S3. Lung tumor analysis and tumor cell proliferation/ death in the $Kras^{G12D/+}$; $p53^{fl/fl}$ model with Keap1/Nrf2 mutation. (A)** Overall tumor burden (%) calculated by dividing the total area of lung tumor by the total area of the lung. *p<0.05 (one-way ANOVA). **(B)** Distribution of tumor number by grade across Keap1/Nrf2 mutant models. N≥9 mice per genotype, *p<0.05 (unpaired t test with Holm-Sidak's multiple comparisons test). $ = fewer than 3 tumors detected across all mice. **(C)** Median tumor size per mouse for each genotype. *p<0.05 (one-way ANOVA). **(D)** Tumor size by grade across all mice per genotype. N≥9 mice and ≥1,900 tumors per genotype. *p<0.05 (Kruskal-Wallis test with Dunn's multiple comparisons test). $ = fewer than 3 tumors detected across all mice. **(E)** Representative immunohistochemical (HC) staining of Ki-67 (scale bars = 20 μM). **(F)** Proportion of tumor cells positive for Ki-67 per mouse. *p<0.05 (one-way ANOVA). **(G)** Percentage of Ki-67 positive tumor cells in each tumor grade per genotype. N=5 mice per genotype, >20,000 tumor cells per mouse. *p<0.05 (unpaired t test with Holm-Sidak's multiple comparisons test). $ = fewer than 3 tumors detected across all mice. For **(B)**, **(D)**, and **(G)**, only one grade 5 tumor was found in the $Keap1^{R554Q/R554Q}$ cohort, and therefore was excluded from these analyses. **(H)** Representative immunohistochemical (IHC) staining of cleaved caspase-3 (CC-3) (scale bars = 20 μM). N=5 mice per genotype. Positive control is BL6 mouse spleen 1 hour after irradiation with 7.5 Gy (scale bar = 10 μM). **(I)** Tumor number per mouse in Keap1/Nrf2 mutant models normalized to lung area. *p<0.05 (one-way ANOVA).

A

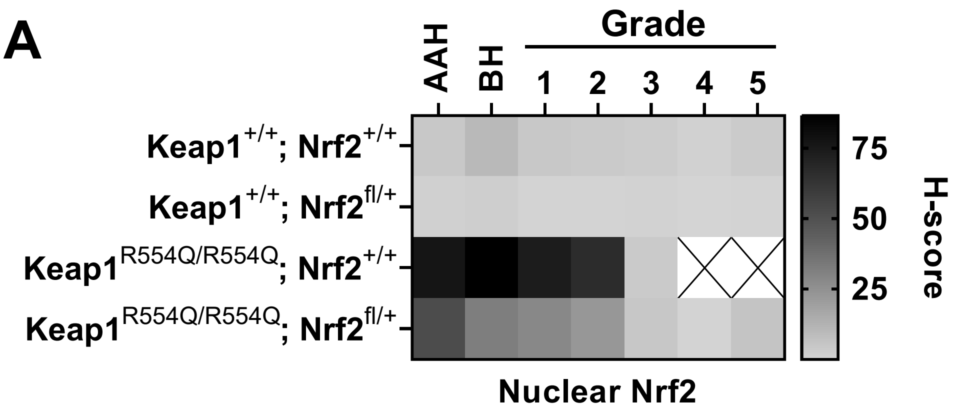

Grade
AAH
BH
1
2
3
4
5
Keap1+/+; Nrf2+/+
Keap1+/+; Nrf2fl/+
Keap1R554Q/R554Q; Nrf2+/+
Keap1R554Q/R554Q; Nrf2fl/+
75
50
25
H-score
Nuclear Nrf2


B

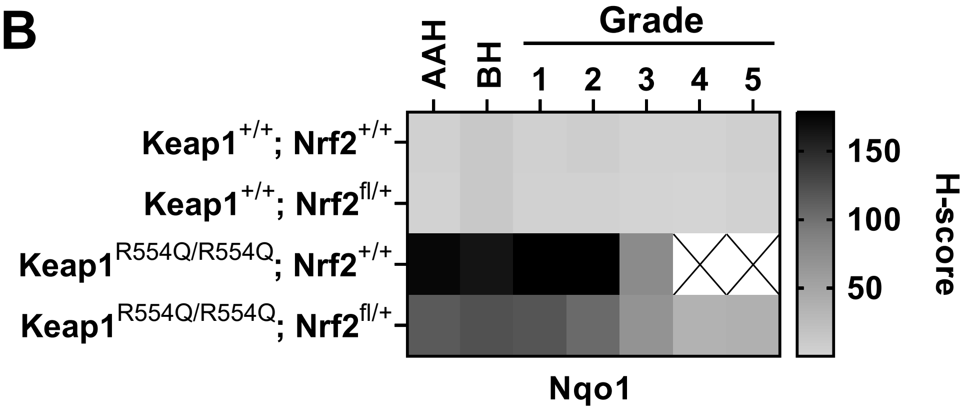

Grade
AAH
BH
1
2
3
4
5
Keap1+/+; Nrf2+/+
Keap1+/+; Nrf2fl/+
Keap1R554Q/R554Q; Nrf2+/+
Keap1R554Q/R554Q; Nrf2fl/+
150
100
50
H-score
Nqo1

**Figure S4. Immunohistochemical analysis of Nrf2 and Nqo1 across tumor grades in the Kras$^{G12D/+}$; p53$^{fl/fl}$ model with single copy Nrf2 deletion. (A, B)** Heatmaps depicting the H-scores per grade from IHC staining for Nrf2 (nuclear) **(A)** and the Nrf2 target Nqo1 (whole cell) **(B)**. N=3 mice per genotype, >20,000 tumor cells per mouse. Only one grade 4 and one grade 5 tumor were found in the Keap1$^{R554Q/R554Q}$; Nrf2$^{+/+}$ cohort, and therefore were excluded from these analyses.

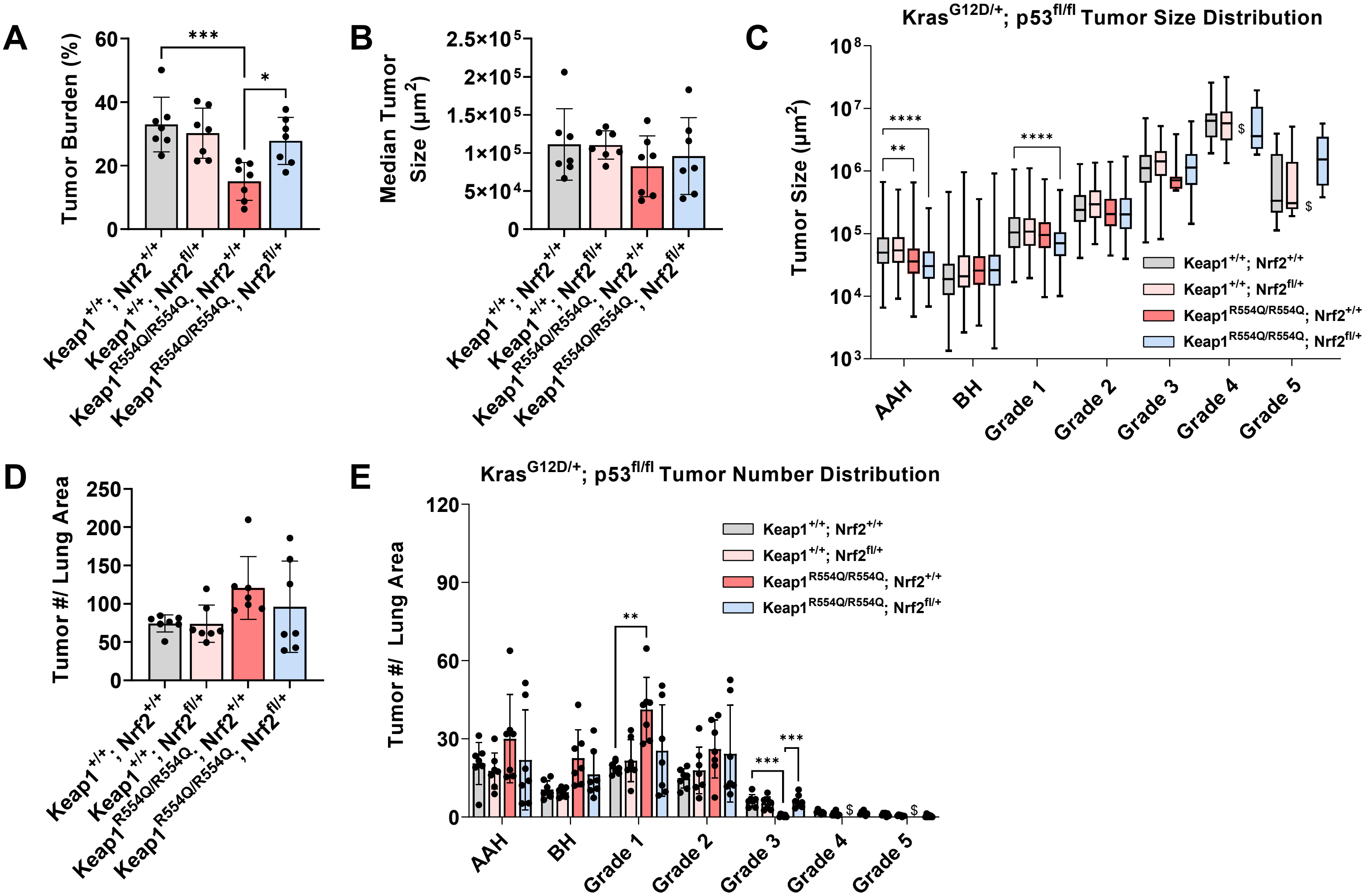
A
Tumor Burden (%)
B
Median Tumor Size (μm2)
C
KrasG12D/+; p53fl/fl Tumor Size Distribution
Tumor Size (μm2)
AAH
BH
Grade 1
Grade 2
Grade 3
Grade 4
Grade 5
Keap1+/+; Nrf2+/+
Keap1+/+; Nrf2fl/+
Keap1R554Q/R554Q; Nrf2+/+
Keap1R554Q/R554Q; Nrf2fl/+
D
Tumor #/ Lung Area
E
KrasG12D/+; p53fl/fl Tumor Number Distribution
Tumor #/ Lung Area

**Figure S5. Lung tumor burden, size, and number quantification in the Kras$^{G12D/+}$; p53$^{fl/fl}$ model with single copy Nrf2 deletion. (A)** Overall tumor burden (%) calculated by dividing the total area of lung tumor by the total area of the lung. **(B)** Median tumor size per mouse for each genotype. For **(A, B)**, *$p<0.05$ (one-way ANOVA). **(C)** Tumor size across all mice per genotype. N=7 mice and $\geq$1,000 tumors per genotype. *$p<0.05$ (Kruskal-Wallis test with Dunn's multiple comparisons test). $ = fewer than 3 tumors detected across all mice. **(D)** Tumor number per mouse normalized to lung area. *$p<0.05$ (one-way ANOVA). **(E)** Distribution of tumor number by grade across Keap1/Nrf2 mutant models. N=7 mice per genotype, *$p<0.05$ (unpaired t test with Holm-Sidak's multiple comparisons test). $ = fewer than 3 tumors detected across all mice. For **(C)** and **(E)**, only one grade 4 and two grade 5 tumors were found in the Keap1$^{R554Q/R554Q}$; Nrf2$^{+/+}$ cohort, and therefore were excluded from these analyses.
